# Replay in Human Visual Cortex is Linked to the Formation of Successor Representations and Independent of Consciousness

**DOI:** 10.1101/2022.02.02.478787

**Authors:** Lennart Wittkuhn, Lena M. Krippner, Christoph Koch, Nicolas W. Schuck

## Abstract

Humans automatically infer higher-order relationships between events in the environment from their statistical co-occurrence, often without conscious awareness. Neural replay of task representations is a candidate mechanism by which the brain learns such relational information or samples from a learned model in the service of adaptive behavior. Here, we tested whether cortical reactivation is related to learning higher-order sequential relationships without consciousness. Human participants viewed sequences of images that followed probabilistic transitions determined by ring-like graph structures. Behavioral modeling revealed that participants acquired multi-step transition knowledge through gradual updating of an internal successor representation (SR) model, although half of participants did not indicate conscious knowledge about the sequential task structure. To investigate neural replay, we analyzed the temporal dynamics of multivariate functional magnetic resonance imaging (fMRI) patterns during brief 10 seconds pauses from the ongoing statistical learning task. We found evidence for backward sequential replay of multi-step sequences in visual cortical areas. These findings indicate that implicit learning of higher-order relationships establishes an internal SR-based map of the task, and is accompanied by cortical on-task replay.

## Introduction

The representation of cognitive maps in the brain is thought to provide the basis for flexible learning, inference, and generalization, and has been a topic of great interest (see e.g., Behrens et al., 2018). While cognitive maps reflect abstract structural knowledge that generalizes specific experiences (Tolman, 1948; Schuck et al., 2018; Behrens et al., 2018), this knowledge must first be learned from individual events that provide structural information only indirectly (Schapiro et al., 2013; Garvert et al., 2017). The brain must therefore extract statistical regularities from continuous experiences, and then use these regularities as the starting point for the formation of abstract, map-like knowledge. A mechanism through which abstract knowledge could then be used to generate flexible behavior is neural replay (Kurth-Nelson et al., 2016; Schuck and Niv, 2019; Liu et al., 2019, 2021; Wittkuhn et al., 2021), the rapid reactivation of sequences simulated from an internal cognitive map (e.g., Wilson and McNaughton, 1994; Foster, 2017). Whether replay only plays a role in *using* cognitive maps, or also in *learning* them, is not known. In the current study, we investigated whether on-task replay of cognitive map-like knowledge plays a role in learning abstract task knowledge from statistical regularities.

The extraction of statistical regularities from experience is known as *statistical learning* (Schapiro and Turk-Browne, 2015; Garvert et al., 2017; Sherman et al., 2020). Statistical learning is automatic and incidental, as it occurs without any instructions or premeditated intention to learn, and often leads to implicit knowledge that is not consciously accessible (Reber, 1989; Seger, 1994; Turk-Browne et al., 2005). In humans, replay from cognitive maps is often studied in participants who have conscious instruction-based task knowledge (e.g., Kurth-Nelson et al., 2016; Liu et al., 2019; Schuck and Niv, 2019). In a statistical learning setting, relationships between events are typically described by pairwise transition probabilities (i.e., the probability that *A* is followed by *B*) to which humans show great sensitivity from an early age on (Saffran et al., 1996). Intriguingly, many experiments have shown that humans extract higher-order relational structures among individual events that go beyond pairwise transition probabilities (for reviews, see e.g., Karuza et al., 2016; Lynn and Bassett, 2020). This includes knowledge about ordinal and hierarchical information that structures individual subsequences (Schuck et al., 2012a,b; Solway et al., 2014; Balaguer et al., 2016), graph topological aspects such as bottlenecks and community structure (Schapiro et al., 2013; Karuza et al., 2017; Kahn et al., 2018), and macro-scale aspects of graph structures (Lynn et al., 2020a,b).

One benefit of abstracted knowledge of transition structures is that it facilitates planing multi-step sequences (Miller and Venditto, 2021; Hunt et al., 2021). While experienced transition structure can be used to learn about the one-step probabilities between pairs of events, it can also be used to compute long-term visitation probabilities, i.e., which events can be expected over a multi-step future horizon. This idea is formalized in the successor representation (SR) model (Dayan, 1993), a predictive map that reflects the (discounted) expected visitations of future events, or states (Garvert et al., 2017; Gershman, 2018; Bellmund et al., 2020; Brunec and Momennejad, 2021; Russek et al., 2021), and can be learned from the experience of individual transitions through a temporal difference (TD) learning mechanism (Dayan, 1993; Russek et al., 2017). The predictive horizon of the SR depends on a discount parameter *γ* which determines how far into the future upcoming states are considered (Momennejad and Howard, 2018; Momennejad, 2020, with 0 *≥ γ ≤* 1; *γ* = 1 indicates the maximum prediction horizon). Based on this previous work, one goal of our study was therefore to investigate whether unconscious statistical learning leads to knowledge of expected future visitations over a predictive horizon, as required for mental simulation.

The second main interest of our study was to understand whether unconscious statistical multi-step knowledge would be reflected in on-task replay. Replay is characterized by the fast sequential reactivation of neural representations that reflect previously experienced transition structure (see e.g., Wikenheiser and Redish, 2015a; Schuck and Niv, 2019; Wittkuhn et al., 2021; Yu et al., 2021). Replay occurs in hippocampal but also cortical brain areas (Ji and Wilson, 2006; Wittkuhn and Schuck, 2021) and has been observed during short pauses from the ongoing task in rodents (Johnson and Redish, 2007; Carr et al., 2011) as well as humans (Schuck and Niv, 2019; Kurth-Nelson et al., 2016; Tambini and Davachi, 2019). Sequential neural reactivation observed during brief pauses from the ongoing task is often referred to as *online* or *on-task* replay. Previous studies have also shown that expectations about upcoming visual stimuli elicit neural signals that are very similar to those during actual perception (Kok et al., 2012, 2014; Hindy et al., 2016; Kok and Turk-Browne, 2018) and anticipatory activation sequences have been found in visual cortex following perceptual sequence learning (Xu et al., 2012; Eagleman and Dragoi, 2012; Gavornik and Bear, 2014; Ekman et al., 2017). It remains unknown, however, whether on-task replay mirrors predictive knowledge that is stored in SR-based cognitive maps, and whether such replay occurs in sensory and motor brain areas. In addition, little is known about to what extent replay is linked to conscious knowledge in humans.

In the present study, we therefore examined whether on-task neural replay in visual and motor cortex reflects anticipation of sequentially structured stimuli in an automatic and incidental statistical learning context. This may reveal if (non-hippocampal) neural replay during on-task pauses interacts with learning of probabilistic cognitive maps. To this end, participants performed an incidental statistical learning paradigm (cf. Schapiro et al., 2012; Lynn et al., 2020a) in which visual presentation order and motor responses followed statistical regularities that were determined by ring-like graph structures. The nature of the graph structure allowed us to dissociate knowledge about individual transition probabilities from an SR-based cognitive map that entails long-term visitation probabilities. Moreover, the transition probabilities among the task stimuli changed halfway through the experiment without prior announcement, which allowed us to understand the dynamical updating of task knowledge and replay within the same participants.

## Results

Thirty-nine human participants took part in an fMRI experiment over two sessions (for details on the study procedure, see Methods and Fig. S1). Participants were first informed that the experiment involves six images of animals (cf. Snodgrass and Vanderwart, 1980; Rossion and Pourtois, 2004, see Fig. S2) and six response buttons mapped onto their index, middle, and ring fingers of both hands (see Fig. S3b). Participants then began the first session of magnetic resonance imaging (MRI), during which they learned the stimulus-response (S-R) mappings between images and response buttons through feedback (*single trials*, Fig. 1a, 8 runs with 60 trials each, 480 trials in total). In single trials, images were shown without any particular sequential order, i.e., all pairwise sequential orderings of the images were presented equally often per run. Participants had to press the button associated with the briefly presented image (500 milliseconds (ms)) during an 800 ms response window (jittered stimulus-response interval (SRI) of 2500 ms on average). While feedback about the correct button was provided on incorrect trials (500 ms), no feedback was given on correct trials. The trial ended with a jittered inter-trial interval (ITI) of 2500 ms on average.

**Figure 1:**
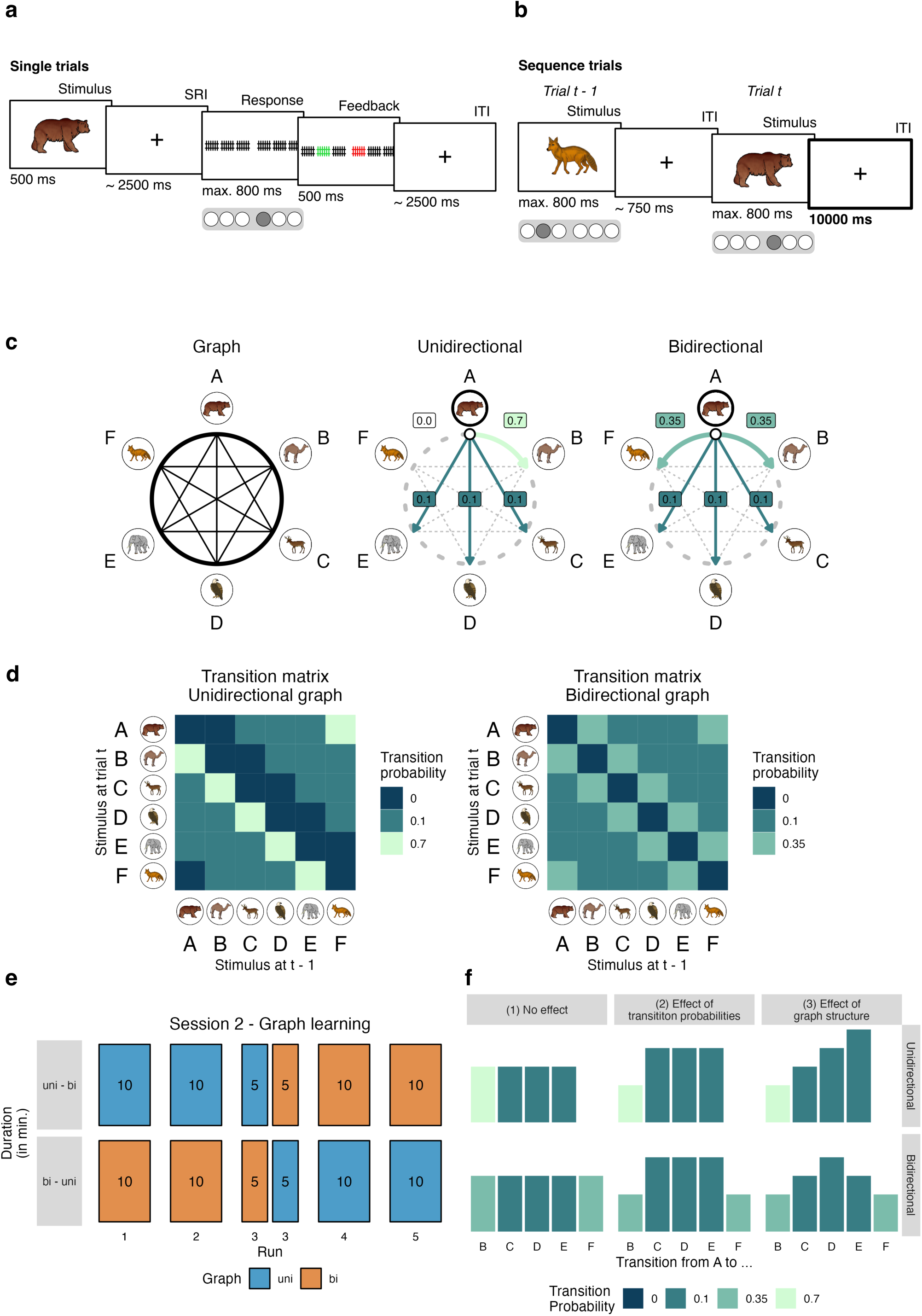
Task design. **(a)** On single trials, individual images were presented for 500 ms. Participants were instructed to press the correct response button associated with the stimulus during the response interval (time limit of 800 ms). Stimulus presentations and motor responses were separated by stimulus-response intervals (SRIs) and inter-trial intervals (ITIs) which both lasted 2.5 s on average (cf. Wittkuhn and Schuck, 2021). Feedback was only presented on incorrect trials (as shown here). Multivariate pattern classifiers were trained on fMRI data from correct single trials only. **(b)** On sequence trials, images were presented for 800 ms, separated by only 750 ms ITIs on average. Participants were asked to press the correct response button associated with the presented stimulus as quickly and accurately as possible within the 800 ms response window. On 10% of trials, ITIs lasted 10 s (*interval trials*; see ITI in trial *t*, highlighted by the thick black border, for illustrative purposes only; 10 s roughly correspond to 8 repetition times (TRs) at a TR of 1.25 s; for details, see Methods). Multivariate pattern classifiers trained on fMRI data from correct single trials were applied to the eight TRs of the 10 s ITIs in sequence trials to investigate task-related neural (re-)activation patterns during on-task intervals. **(c)** The relationships among the six task stimuli depicted as a ring-like graph structure (left). In the unidirectional graph (“uni”; middle), stimuli frequently transitioned to the clockwise neighboring node (*p_ij_* = *p_AB_* = 0.7), never to the counterclockwise neighboring node (*p_AF_* = 0.0), and only occasionally to the three other nodes (*p_AC_* = *p_AD_* = *p_AE_* = 0.1). In the bidirectional graph (“bi”; right), stimuli were equally likely to transition to the clockwise or counterclockwise neighboring node (*p_AB_* = *p_AF_* = 0.35) and only occasionally transitioned to the three other nodes (*p_AC_* = *p_AD_* = *p_AE_* = 0.1). Transition probabilities are highlighted for node *A* only, but apply equally to all other nodes. Arrows indicate possible transitions, colors indicate transition probabilities (for a legend, see panel 1d). **(d)** Transition matrices of the unidirectional (left) and bidirectional (right) graph structures. Each transition matrix depicts the probability (colors; see legend) of transitioning from the stimulus at the previous trial *t −* 1 (x-axis) to the current stimulus at trial *t* (y-axis). **(e)** Within-participant order of the two graph structures across the five runs of the sequence trials. *n* = 12 participants first experienced the unidirectional, then the bidirectional graph structure (uni – bi; top horizontal panel) while *n* = 27 participants experienced the reverse order (bi – uni; bottom horizontal panel). In both groups of participants, the graph structure was changed without prior announcement halfway through the third task run. Numbers indicate approximate run duration in minutes (min). Colors indicate graph condition (uni vs. bi; see legend). **(f)** Visualization of the relative magnitude of the outcome variable (e.g., behavioral responses or classifier probabilities; y-axis) for specific transitions between the nodes (x-axis) and the two graph structures (uni vs. bi; horizontal panels) under the three assumptions (vertical panels), (1) that there is no difference between transitions (null hypothesis), (2) that outcome variables are only influenced by the one-step transition probabilities between the nodes (colors), or (3) that outcome variables are influenced by the multi-step relationships between nodes in the graph structure (or the distance between nodes in the graph structure, here referred to as *node distance*). An effect of unidirectional graph structure would be evident in a linear relationship between node distance and the outcome variable, whereas a bidirectional graph structure would be reflected in a U-shaped relationship between node distance and outcome measures (possibly inverted, depending on the measure). The stimulus material (individual images of a bear, a dromedary, a dear, an eagle, an elephant and a fox) shown in (a), (b), and (c) were taken from a set of colored and shaded images commissioned by Rossion and Pourtois (2004), which are loosely based on images from the original Snodgrass and Vanderwart set (Snodgrass and Vanderwart, 1980) (for an overview of all images, see Fig. S2). The images are freely available from the internet at https://sites.google.com/andrew.cmu.edu/tarrlab/stimuli under the terms of the Creative Commons Attribution-NonCommercial-ShareAlike 3.0 Unported license (CC BY-NC-SA 3.0; for details, see https://creativecommons.org/licenses/by-nc-sa/3.0/). Stimulus images courtesy of Michael J. Tarr, Carnegie Mellon University (for details, see http://www.tarrlab.org/).

The second session started with one additional run of single trials that was followed by five runs of *sequence trials* (Fig. 1b, 240 trials per run, 1200 trials in total). As before, participants had to press the correct button in response to each animal image. Participants were only informed that in this task condition no feedback would be provided and images would be shown at a faster pace (800 ms per image and 750 ms ITI between images on average), with occasional 10 seconds (s) breaks in between (10% of trials, i.e., 120 of all trials per participant during which only a fixation cross was on the screen, henceforth *interval trials*; 24 trials per class). Unbeknownst to the participants, the images also started to follow a probabilistic transition structure (for details, see below and Fig. 1c–d). At the end of the second session, participants completed a post-task questionnaire assessing explicit sequence knowledge.

The sequential ordering of images during sequence trials was determined by either a unidirectional or bidirectional ring-like graph structure with probabilistic transitions (Fig. 1c–d; for details, see Methods). In the unidirectional graph condition (Fig. 1c, middle, henceforth *uni*), each image had one frequent transition to the clockwise neighboring node (probability of *p_ij_* = 0.7), never transitioned to the counterclockwise neighbor (*p_ij_* = 0.0), and was followed occasionally by the three other nodes (*p_ij_* = 0.1 each; Fig. 1d, left). In consequence, stimuli most commonly transitioned in clockwise order along the ring shown in Fig. 1c. In the bidirectional graph condition (Fig. 1c, right, henceforth *bi*), transitions to both neighboring nodes (clockwise and counterclockwise) were equally likely (*p_ij_* = 0.35), and transitions to all other three nodes occurred with *p_ij_* = 0.1 (Fig. 1d, right), as in the unidirectional graph. Every participant started the task in one of these conditions (uni or bi). Halfway through the third run, transitions began to be governed by the alternative graph, such that all participants experienced both graphs as well as the change between them (Fig. 1e). 12 participants started in the unidirectional condition and transitioned to the bidirectional graph (uni – bi; top horizontal panel in Fig. 1e), while 27 participants experienced the reverse order (bi – uni; bottom horizontal panel in Fig. 1e).

### Behavioral results

We first asked whether participants learned the stimulus-response (S-R) mappings between animal images and response keys sufficiently well. Behavioral accuracy on single trials (Fig. 1a) surpassed the chance-level (16.67%) in all runs (*x̄ ≥* 86.50%, confidence intervals (CIs) [*≥* 80.79, +*∞*], *t*_38_ *≥* 20.62, *p*s *<* 0.001 (corrected), *d*s *≥* 3.30; Figs. 2a, S4a). The same was true during sequence trials (*x̄ ≥* 85.12, CIs [*≥* 82.55, +*∞*], *t*_38_ *≥* 44.90, *p*s *<* 0.001 (corrected), *d*s *≥* 7.19; Figs. 1b, 2a, S4b), where behavioral accuracy also improved with time (effect of run: *F*_1.00,38.00_ = 7.96, *p* = 0.008, Fig. S4b).

**Figure 2:**
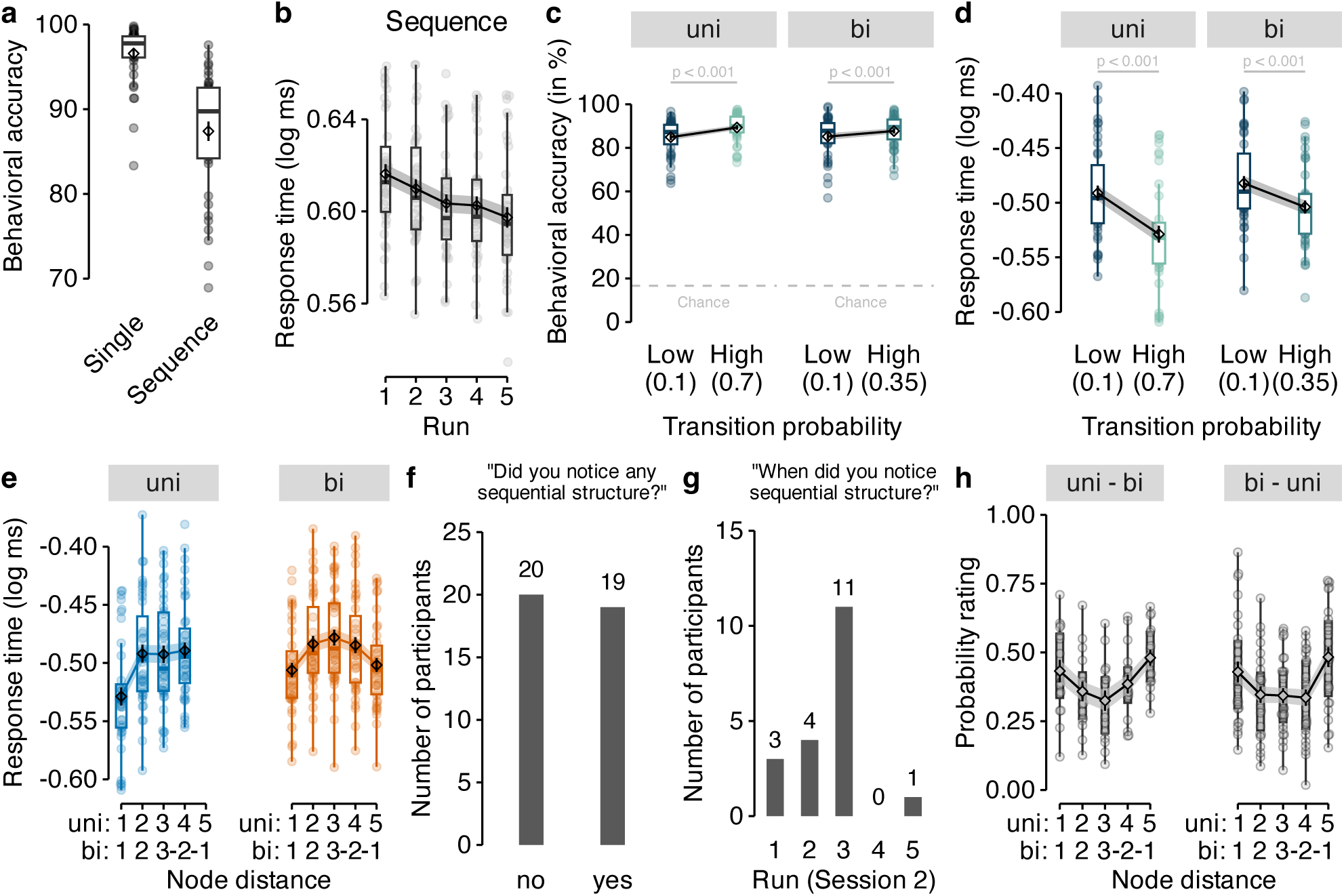
Behavioral responses are influenced by transition probabilities and graph structure. **(a)** Mean behavioral accuracy (in %; y-axis) across all nine runs of the single trials (left) and across all five runs of the sequence trials (right). **(b)** Mean log response time (y-axis) per run (x-axis) in sequence trials. **(c)** Mean behavioral accuracy (y-axis) following transitions with low (*p_ij_* = 0.1) and high probability (x-axis; *p_ij_* = 0.7 and *p_ij_* = 0.35 in the uni- and bidirectional graph conditions of sequence trials, respectively) for both graph structures (panels). Colors as in Figs. 1c and 1d. The horizontal dashed lines indicate the chance level (16.67%). **(d)** Mean log response time (y-axis) following transitions with low (*p_ij_* = 0.1) and high probability (*p_ij_* = 0.7 and *p_ij_* = 0.35 in the uni- and bidirectional graph conditions of sequence trials, respectively) for both graph structures (panels). Colors as in panel (c), Figs. 1c and 1d.**(e)** Mean log response time (y-axis) as a function of uni- or bidirectional node distance (x-axis) in data from the two graph structures (colors / panels). Colors as in Fig. 1e. **(f)** Number of participants (y-axis) indicating whether they had noticed any sequential ordering during the graph task (“no” or “yes”; x-axis). **(g)** Number of participants (y-axis) who had detected sequential ordering (*n* = 19, see panel (f)) indicating in which of the five runs of the sequence task (x-axis) they had first noticed sequential ordering. **(h)** Ratings of pairwise transition probabilities (in %; y-axis) as a function of node distance, separately for both orders of graph structure (uni – bi vs. bi – uni; panels). Boxplots in (a), (b), (c), (d), (e) and (h) indicate the median and interquartile range (IQR). The lower and upper hinges correspond to the first and third quartiles (the 25^th^ and 75^th^ percentiles). The upper whisker extends from the hinge to the largest value no further than 1.5*∗* IQR from the hinge (where IQR is the interquartile range (IQR), or distance between the first and third quartiles). The lower whisker extends from the hinge to the smallest value at most 1.5*∗* IQR of the hinge. The diamond shapes in (a), (b), (c), (d), (e) and (h) show the sample mean. Error bars and shaded areas in (a), (b), (c), (d), (e) and (h) indicate *±*1 standard error of the mean (SEM). Each dot in (a), (b), (c), (d), (e) and (h) corresponds to averaged data from one participant. All statistics have been derived from data of *n* = 39 human participants who participated in one experiment.

Next, we investigated sequential knowledge. Although participants were not informed that images during sequence trials followed a sequential structure, we expected that incidental learning would allow them to anticipate upcoming stimuli during these trials, and thus respond faster with learning. A linear mixed effects (LME) model that tested the effect of task run on response times was in line with this idea, showing a significant decrease of response times over the course of learning, *F*_1.00,38.00_ = 25.86, *p <* 0.001 (Fig. 2b). More directly, we expected that participants would learn the probabilistic transition structure during sequence trials, and the change therein in middle of the third run. Indeed, participants responded faster and more accurately to one-step transitions with high compared to low probabilities in the unidirectional graph condition (*p_ij_* = 0.7 (high) vs. *p_ij_* = 0.1 (low) transition probabilities, *p*s *<* 0.001), and in the bidirectional graph condition (*p_ij_* = 0.35 (high) vs. *p_ij_* = 0.1 (low), all *p*s *<* 0.001 (corrected), *d*s *≥* 0.67; Figs. 2c–d).

Our main behavioral hypothesis was that participants would not only learn about one-step transition probabilities, but also form internal maps of the underlying graphs that reflect the higher-order structure of statistical multi-step relationships between stimuli, i.e., how likely a particular stimulus will be experienced in two, three, or more steps from the current time point (cf. Lynn and Bassett, 2020; Lynn et al., 2020a). In our task, this meant that participants might react differently to the three transitions that all have the same one-step transition probability, since they differ in how likely they would occur in multi-step trajectories. For instance, the one-step transition probabilities for A*→*C, A*→*D, and A*→*E were the same in the unidirectional graph (*p_ij_* = 0.1, see the graph structures in Fig. 1c), but the two-step probability of A*→*C was higher than for the other transitions, since the most likely two-step path was A*→*B*→*C, implying that participants should react faster to A*→*C than to A*→*D transitions. For simplicity, we will henceforth refer to the A*→*C transition as having a shorter “node distance”, than A*→*D or A*→*E (see the rightmost column in Fig. 1f, where colors reflect one-step transition probabilities, and the height of the bars indicate node distance). Analyzing response times as a function of the node distance (Fig. 1f; for details, see Methods) indicated a significant effect of node distance on response times in both unidirectional, *F*_1.00,115.78_ = 44.34, *p <* 0.001, and bidirectional data, *F*_1.00,38.00_ = 57.36, *p <* 0.001 (Fig. 2e).

We assessed whether participants were able to express knowledge of the sequential ordering of stimuli and graph structures explicitly during a post-task questionnaire. Asked whether they had noticed any sequential ordering of the stimuli in the preceding sequence task, *n* = 19 participants replied “yes” and *n* = 20 replied “no” (Fig. 2f). Performance on the sequence task did not differ significantly between participants with and without conscious knowledge, as the effect of node distance on response times (see above) was comparable between these two groups (*p*s *≥* 0.48 for each node distance). Of those participants who noticed sequential ordering (*n* = 19), almost all (18 out of 19) indicated that they had noticed ordering within the first three runs of the task (Fig. 2g), and more than half of those participants (11 out of 19) indicated that they had noticed ordering during the third task run, during which the graph structure was changed. Thus, sequential ordering of task stimuli remained at least partially implicit in half of the sample, and the change in the sequential order halfway through the third run of graph trials seemed to be one potential cause for the conscious realization of sequential structure. Asked to rate the transition probabilities of all pairwise sequential combinations of the six task stimuli (30 ratings in total), participants reported probability ratings that reflected the bidirectional graph structure on average (Fig. 2h), i.e., probabilities of clockwise and counterclockwise transitions were rated higher than rarer transitions to intermediate nodes (regardless of the order in which participants had experienced the two graph structures immediately before the questionnaire).

To investigate how multi-step knowledge emerged from experience, we next asked whether response times were influenced by the *n*-step probabilities that participants had experienced up until each trial. Specifically, we used a successor representation (SR) model (Dayan, 1993) that iteratively learns the discounted long-term occupation probabilities of every node starting from all other nodes. In this model, each node is associated with a vector that reflects the probability that starting from the current node a participant would experience any of the other nodes over a future-discounted predictive horizon. While these SR-vectors were uniformly initialized at the beginning of the task, they were dynamically updated following each transition experienced in the task, using a temporal difference (TD) learning rule (Dayan, 1993; Russek et al., 2017). Concretely, after experiencing the transition from image *s_t_* to *s_t_*_+1_, the row corresponding to image *s_t_* of the successor matrix **M** was updated as follows

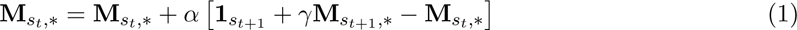

whereby **1***_st_*_+1_ is a one-hot vector with a 1 in the *s_t_*_+1_^th^ position, and *α* (“alpha”) is a learning rate. The discounting parameter *γ* (“gamma”) defines the extent to which multi-step transitions are taken into account, which we will henceforth refer to as the “predictive horizon” (cf. Gershman et al., 2012; Momennejad, 2020).

To test whether this model accounted for the emergence of multi-step knowledge in our task, we first computed a series of SR models that covered the continuum between mere one-step learning (*γ* = 0) and learning over a large predictive horizon (*γ* = 0.95, models in steps of 0.05), using the exact stimulus sequences that each participant experienced in the task. We then asked how well response times were predicted by the myopic compared to the more far-sighted SR models by comparing the Akaike information criterion (AIC) scores of corresponding LME models. Each LME model regressed one participant’s trial-by-trial development of multi-step knowledge as predicted by the SR model against their trial-by-trial response times, wherein transitions that were less likely according to the model should be associated with longer response times (the successor matrix **M** was converted to a Shannon surprise predictor (cf. Shannon, 1948), and LME models included fixed effects of task run, graph and graph order, and by-participant random intercepts and slopes; for details, see Methods). This analysis showed that a discount parameter of *γ* = 0.3 resulted in the lowest AIC score (Fig. 3a), and models with non-zero *γ* parameters yielded substantially better fits than a model which assumed only knowledge of one-step transitions (*γ* = 0, leftmost data point in Fig. 3a). Thus, participants’ response times clearly indicated multi-step graph knowledge consistent with SR models. Separate analyses for the two graph structures (uni vs. bi) and graph orders (uni – bi vs. bi – uni), showed that non-zero *γ* parameters achieved better fits in all cases, and indicated some differential effects of the *γ* parameter depending on graph structure and graph order (Fig. S5).

**Figure 3:**
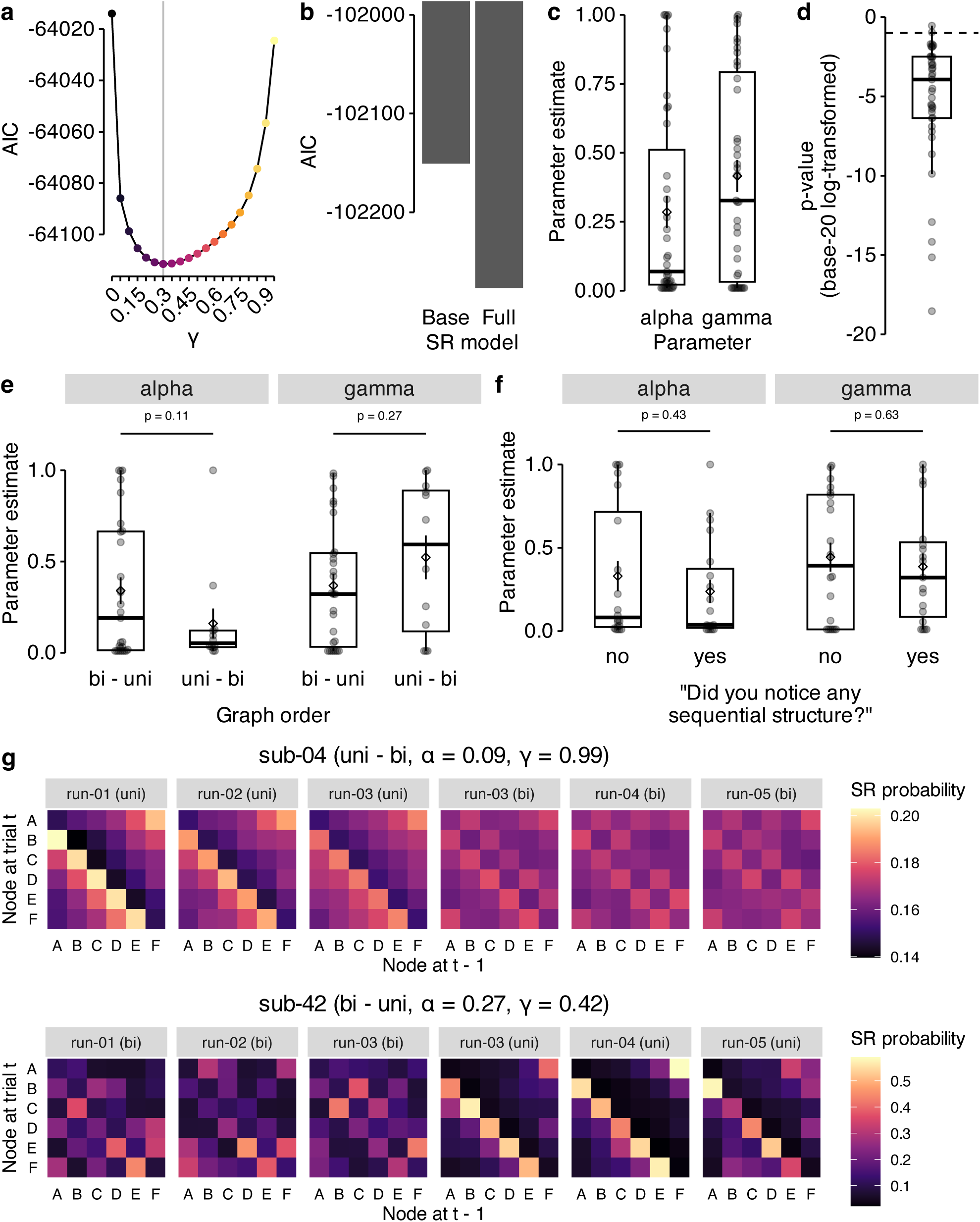
Successor representation (SR) modeling of behavior. **(a)** AIC scores (y-axis) for LME models fit to participants’ log response time data using Shannon surprise based on SRs with varying predictive horizons (the discounting parameter *γ*; x-axis) as the main predictor variable. The vertical line marks the lowest AIC score (at *γ* = 0.3). **(b)** AIC scores (y-axis) for baseline SR model where the *γ* parameter was fixed to 0 and only the *α* parameter was fit per participant (left bar) compared to the full SR model where both the *γ* and *α* parameter were fit to individual participants’ data (right bar). AIC scores were adjusted for the number of free parameters (one free parameter for the baseline and two free parameters for the full SR model). **(c)** Model parameter estimates for *α* and *γ* in the full _SR_ model after participant-specific model fitting. **(d)** *p*-values (on *log*_20_ scale) of the fixed effect of the SR-based Shannon surprise in the general linear model (GLM) that, in combination with the SR-model parameters *α* and *γ*, best explained participants’ response times. The dashed line indicates an alpha level of 0.05 (*log*_20_(0.05) = 1) with lower values indicating *p <* 0.05. **(e)** Parameter estimates as in panel (c) and (f) but separated by the graph order that the participant experienced (uni–bi vs. bi–uni). **(f)** Parameter estimates as in panel (c) and (e) but separated by conscious knowledge (“yes” vs. “no”). **(g)** Successor representation (SR) matrices for two selected participants who experienced the two graph structures in uni - bi (top panel) or bi - uni (bottom panel) order, respectively, separately for each combination of run and graph structure in sequence trials (panels) at the last trial of the respective task section. The colors indicate the normalized expected future visitation of each of the six nodes in the graph structure according to Equation 2. SR matrices were determined based on individually fitted parameters for *α* and *γ* (see plot titles). Note, that the third run included the change from one to the other graph structure and the data is therefore shown separately for the two halves of the run. Boxplots in (c), (d), (e), and (f) indicate the median and IQR. The lower and upper hinges correspond to the first and third quartiles (the 25^th^ and 75^th^ percentiles). The upper whisker extends from the hinge to the largest value no further than 1.5*∗* IQR from the hinge (where IQR is the interquartile range (IQR), or distance between the first and third quartiles). The lower whisker extends from the hinge to the smallest value at most 1.5*∗* IQR of the hinge. The diamond shapes in (c), (d), (e), and (f) show the sample mean. Error bars in (c), (d), (e), and (f) indicate *±*1 SEM. Each dot in (c), (d), (e), and (f) corresponds to averaged data from one participant. All statistics have been derived from data of *n* = 39 human participants who participated in one experiment.

In order to get an estimate of multi-step learning per participant, we fitted the *γ* and *α* parameters of the SR model to each participant’s data (for details, see Methods). In line with the above analysis performed across all participants’ data, we found an improvement in fit when the model had a free γ parameter for each participant, compared to a baseline model in which *γ* was fixed to zero (Fig. 3b; AICs: *−*99789.02 vs. *−*99668.43 for baseline model; smaller values indicate better fit). 27 out of 39 participants (about 70%) had estimates of *γ >* 0.1, and the mean of the best fitting parameters was *γ* = 0.41 (standard deviation (SD): *σ*(*γ*) = 0.36; Fig. 3c), indicating multi-step graph knowledge consistent with SR models. The average learning rate was *α* = 0.28 (SD: *σ*(*α*) = 0.35; Fig. 3c). As a sanity check, we also verified that the Shannon surprise predictor derived from the individually fitted model parameters actually had a significant effect on participants’ response times, which was the case for almost all participants (37 of 39, i.e., ca. 95%; Fig. 3d; using an *α*-level of 0.05). Splitting the data by graph order (uni–bi vs. bi–uni) did not result in significant differences in parameter estimates for both *α* and *γ* (*t*s *≥* 1.65, *p*s *≥* 0.11, *d*s *≥* 0.14; Fig. 3e). Conscious knowledge (“yes” vs. “no”) was not related to significant differences in parameter estimates for both *α* and *γ* (*t*s *≥* 0.49, *p*s *≥* 0.43, *d*s *≥* 0.16; Fig. 3f). Fitted SR matrices for two example participants are shown in Fig. 3g. For illustrative purposes, we selected one participant with a deep predictive horizon (*γ* = 0.99) and a participant with fitted parameters close to the mean parameters in the sample (*α* = 0.27, *γ* = 0.42). The SR matrices of all participants can be found in Figs. S6, S7.

### fMRI results

To ask whether learning of map-like graph representations was accompanied by on-task neural replay, we first trained fMRI pattern classifiers that could detect stimulus-related activity. Logistic regression classifiers were trained on fMRI signals related to stimulus and response onsets in correct single trials, where category order was random and trials were sufficiently separated by stimulus-response intervals (SRIs) and inter-trial intervals (ITIs) of 2500 ms each (one TR per trial; onsets shifted by 4 s after stimulus or motor onset; one-versus-rest training; for details, see Methods; cf. Wittkuhn and Schuck, 2021). Separate classifiers were trained on data from gray-matter-restricted anatomical regions of interest (ROIs) of occipito-temporal cortex and pre- and postcentral gyri, which reflect visual object processing (cf. Haxby et al., 2001) and sensorimotor activity (e.g., Kolasinski et al., 2016), respectively. The trained classifiers successfully distinguished between the six animals on single trials. Leave-one-run-out classification accuracy was *M* = 63.08% in occipito-temporal data (*SD* = 12.57, *t*_38_ = 23.06, CI [59.69, +*∞*], *p <* 0.001, compared to chance level of 16.67%, *d* = 3.69) and *M* = 47.05% in motor cortex data (*SD* = 7.79%, *t*_38_ = 24.36, CI [44.95, +*∞*], *p <* 0.001 vs. chance, *d* = 3.90, all *p*-values Bonferroni-corrected, Fig. 4a). Training only on data from session 1 (eight runs of single trials) and testing on data from session 2 (one run of single trials) indicated no decoding decrements compared to within-session testing, *F*_8_,_655_ = 0.95, *p* = 0.48 (Fig. S8; for details, see Methods). A time-resolved analysis of classifier probabilities over fifteen time points (TRs of 1.25 s) following event onsets showed that the normalized probability of the true stimulus class given the data peaked at the fourth TR (3.75 to 5.0 s; Fig. 4b) as expected based on our previous work (Wittkuhn and Schuck, 2021). During this peak TR, the probability of the true class (i.e., the class of the current trial) was significantly higher than the mean probability of all other classes (difference between current vs. other animals in visual ROI: *M* = 17.88, *t*_38_ = 21.72, CI [16.22, 19.55], *p <* 0.001, *d* = 3.48; motor ROI: *M* = 12.24, *t*_38_ = 32.10, CI [11.47, 13.01], *p <* 0.001, *d* = 5.14, all *p*-values Bonferroni-corrected; Fig. 4b). We next applied the trained classifiers to data from the sequence task, where on some trials participants experienced an extended post-stimulus interval of 10 s (roughly 8 TRs) during which only a fixation cross was displayed (120 trials per participant in total; 24 trials per class). As expected, the classifier probability of the animal displayed at the beginning of the extended post-stimulus interval (or the corresponding motor response, respectively) was higher compared to all other classes (Fig. 4c), and rising and falling slowly as observed in single trials (Fig. 4d; mean probability of current event vs. all others in both ROIs; *t*s *≥* 11.92, *p*s *< .*001, *d*s *≥* 1.91, *p*-values Bonferroni-corrected).

**Figure 4:**
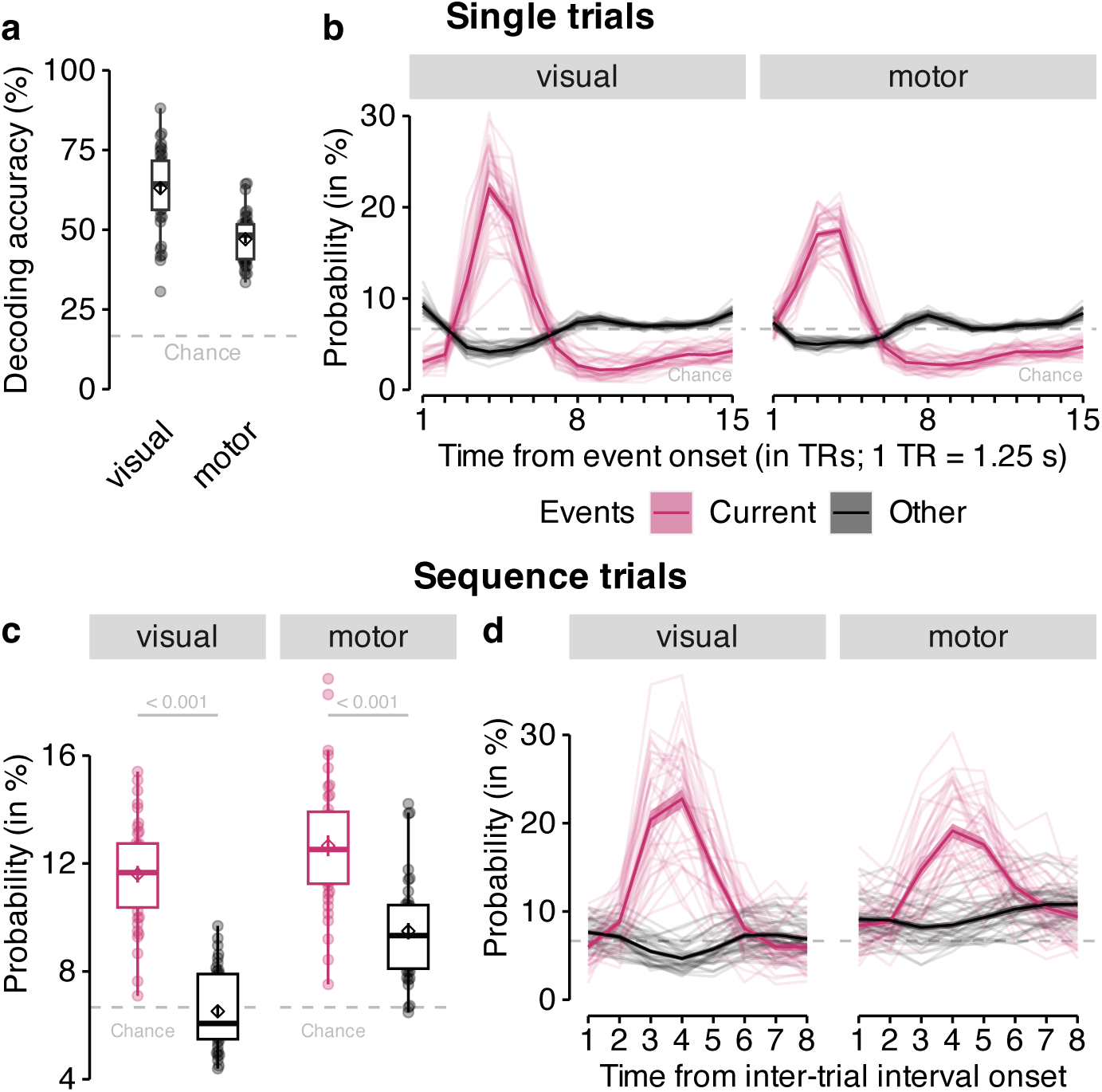
Classification accuracy and probabilistic classifier time courses on single and sequence trials. **(a)** Cross-validated leave-one-run-out classification accuracy (in %; y-axis) in decoding six unique visual objects in occipitotemporal data (“visual”; left) and six unique motor responses in sensorimotor cortex data (“motor”; right) during task performance on single trials. Chance level is at 16.67% (horizontal dashed line). **(b)** Time courses (in TRs from stimulus / response onset; x-axis) of probabilistic classification evidence (in %; y-axis) for the event on the current single trial (purple color) compared to all other events (black color), separately for both ROIs (panels). Classifier probabilities in single trials were normalized across 15 TRs. The chance level therefore is at 100*/*15 = 6.67% (horizontal dashed line). **(c)** Mean classifier probability (in %; y-axis) for the event that occurred on the current sequence trial (purple color), shortly before the onset of the on-task interval, compared to all other events (black color), averaged across all eight TRs in the on-task interval, separately for each ROI (panels). **(d)** Time courses (in TRs from on-task interval onset; x-axis) of mean probabilistic classification evidence (in %; y-axis) in sequence trials for the event that occurred on the current trial (purple color) and all other events (back color). Classifier probabilities in (c) and (d) were normalized across 8 TRs. The chance level therefore is at 100*/*8 = 12.5% (horizontal dashed line). Boxplots in (a) and (c) indicate the median and IQR. The lower and upper hinges correspond to the first and third quartiles (the 25*^th^* and 75*^th^* percentiles). The upper whisker extends from the hinge to the largest value no further than 1.5*∗* IQR from the hinge (where IQR is the interquartile range (IQR), or distance between the first and third quartiles). The lower whisker extends from the hinge to the smallest value at most 1.5*∗* IQR of the hinge. The diamond shapes in (a) and (c) show the sample mean. Error bars and shaded areas indicate *±*1 SEM. Each dot in (a) and (c) or line in (b) and (d) corresponds to averaged data from one participant. 1 TR = 1.25 s. All statistics have been derived from data of *n* = 39 human participants who participated in one experiment.

Because our main focus was on replay rather than stimulus-driven activity, we removed classifier probabilities of the current stimulus in interval trials from all following analyses and asked whether we could detect sequential replay of the five stimulus categories that had *not* been displayed on the current trial. As in our previous work, we looked for evidence of replay in the ordering of classification probabilities within single TRs (Wittkuhn and Schuck, 2021), expecting that this metric would reflect reactivation of either past events or anticipated upcoming events (backward or forward replay). A specific question of our work was whether replay would involve only one-step transitions, or multi-step transitions derived from each participant’s best fitting SR model, as described above. Qualitatively, a one-step perspective predicts that in unidirectional sequence trials the classifier of either stimulus *B* or *F* should be activated following the presentation of *A*, given that both were one step away from image *A* in the forward and backward directions (see Fig. 1c). The additional expectation derived from the SR-model was that, although images *C*, *D*, and *E* had equal one-step forward transition probabilities, the corresponding classifier probabilities could reflect their multi-step SR-probabilities (C *>* D *>* E), with the same logic applying to backward transitions. Following our previous work (Wittkuhn and Schuck, 2021), we also assumed that the ordering during the earlier phase of the on-task interval (TRs 1–4) would reflect the true directionality of the replayed sequence and would be reversed in the later phase of the interval (TRs 5–8), reflecting the rising and falling slopes of the underlying hemodynamic response functions (HRFs). For example, forward replay would be indicated by forward sequentiality in earlier TRs and backward sequentiality in later TRs, while the reverse would be true for backward replay, i.e., backward sequentiality in earlier TRs and forward sequentiality in later TRs.

A major obstacle for replay analyses during brief pauses from ongoing behavior is that the sequential ordering of previously displayed stimuli will lead to residual stimulus-evoked activation that can bias any analysis of sequential reactivation. To account for this, we modeled the stimulus-driven classifier probabilities based on the specific previous trial history of each interval, and asked whether the observed classifier probabilities reflected ordering above and beyond the trial history effects, lever-aging, in particular, variability due to probabilistic transitions. Stimulus-driven classifier effects were modeled based on sine-based response functions that were fit to the time courses in (independent) single trials, as done previously in Wittkuhn and Schuck, 2021 (with parameters for amplitude, response duration, onset delay and baseline; see Methods and Fig. S9 for example fits and comparisons across stimulus categories). The fitted participant- and stimulus-specific response functions were then convolved with the onsets of ten stimuli preceding and following each interval trial (Figs. 5a and S10). We used the resultant predicted time courses of stimulus-driven classifier probabilities to set up a baseline LME model of observed classifier evidence during interval trials.

**Figure 5:**
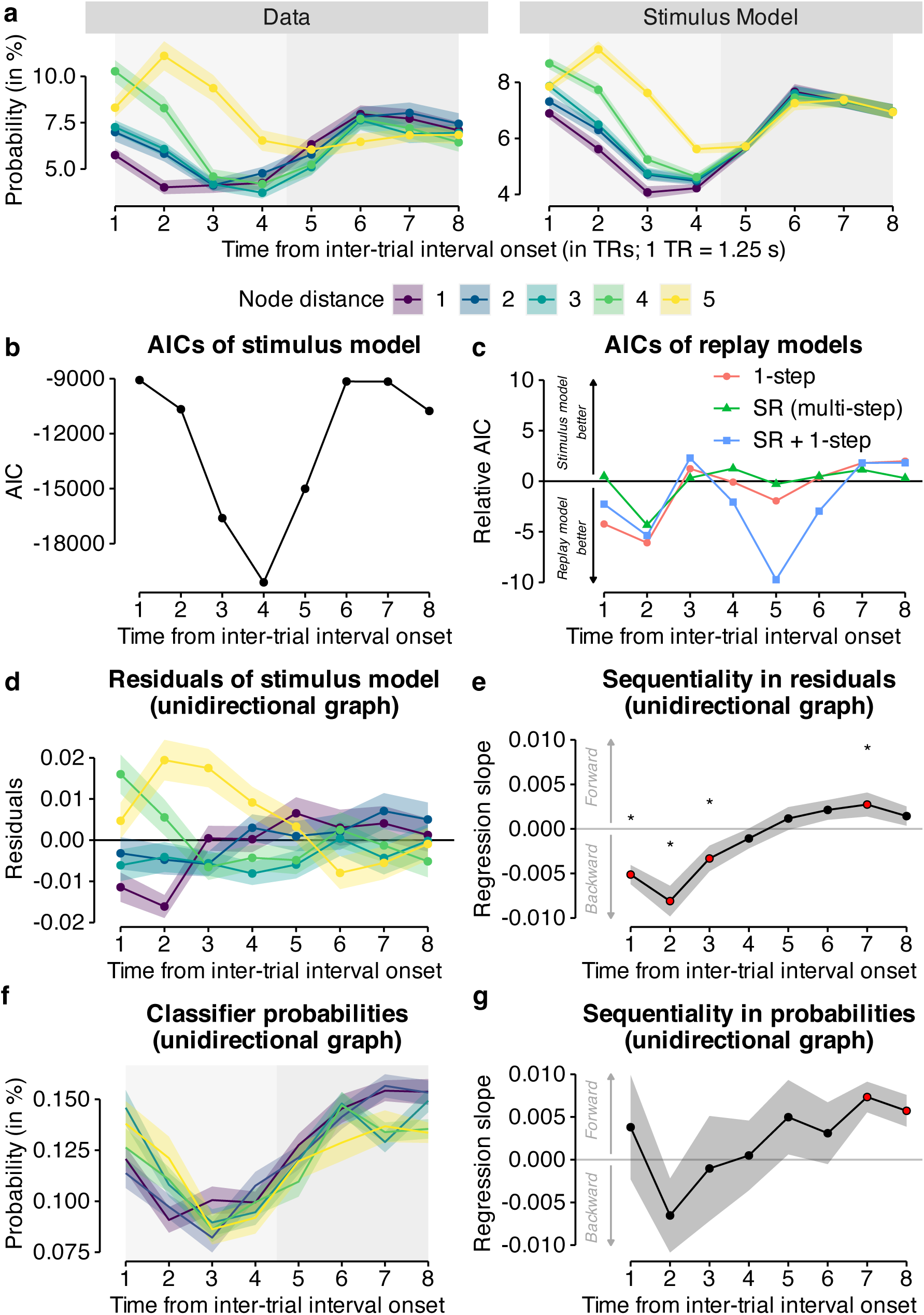
Classifier probabilities during on-task intervals of sequence trials are modulated by one-step and multi-step transition probabilities after accounting for stimulus-evoked activity. **(a)** Time courses (in TRs from inter-trial interval (ITI) onset; x-axis) of observed classifier probabilities (in %; y-axis) in the data from interval trials with unidirectional graph structure (“Data”; left panel) compared to modeled probabilistic classifier evidence (in %; y-axis) based on the sine wave response function with individually fitted parameters (“Stimulus Model”; right panel) separately for the five node distances in unidirectional graph data (colors; see legend). **(b)** Time course (in TRs from ITI onset; x-axis) of AIC scores (y-axis) from the baseline model (including only nuisance regressor for stimulus-evoked activity). **(c)** Time courses (in TRs from ITI onset; x-axis) of relative AIC scores from LME models (y-axis) including predictors for (1) stimulus-evoked activation (baseline model) and, in addition to this baseline model, (2) one-step transition probabilities, (3) multi-step transition probabilities according to the SR model, (4) one-step transition and multi-step SR probabilities. Positive and negative values indicate a worse and better fit, respectively, of the LME model compared to the baseline model that accounts for stimulus-evoked activation but does not include additional predictors. **(c)** Time courses (in TRs from ITI onset; x-axis) of the residuals of the stimulus model separately for the five node distances in unidirectional graph data (colors; see legend in panel a). **(e)** Time courses (in TRs from ITI onset; x-axis) of mean regression slopes (y-axis) relating node distance (i.e., sequential position from current node in the graph structure) to their residuals in the stimulus model as in (d). Positive and negative values indicate forward and backward sequentiality, respectively. Red dots and asterisks indicate significant differences from baseline (horizontal gray line at zero; all *p*s *≤ .*05, uncorrected; two-sided one-sample t-tests, one test per TR). **(f) (f)** Time courses (in TRs from ITI onset; x-axis) of mean probabilistic classification evidence (in %; y-axis) for each of the five classes, colored by node distance (colors; see legend in panel a) for data from the unidirectional graph condition and occipito-temporal anatomical ROIs after removing TRs expected to contain stimulus-evoked activation based on our modeling approach. **(g)** Time courses (in TRs from ITI onset; x-axis) of mean regression slopes (in %) relating node distance to normalized classifier probability (see panel (f)). Positive and negative values indicate forward and backward sequentiality, respectively. Red dots indicate significant differences from baseline (horizontal gray line at zero; all *p*s *≤ .*05, false discovery rate (FDR)-corrected; two-sided one-sample t-tests, one test per TR). Shaded areas in (a), (d), (e), (f) and (g) represent *±*1 SEM. All statistics have been derived from data of *n* = 39 human participants who participated in one experiment. 1 TR = 1.25 s.

Using this approach, we first investigated replay in the visual ROI. As expected, the nuisance regressors reflected the overall activation patterns well (Fig. 5a; left panel), resulting in a good overall fit of the baseline model, peaking at the fourth TR after the last stimulus was shown (*p < .*001, Fig. 5b). To investigate replay, we compared this baseline model to a suite of models that included additional regressors for either one-step task transition probabilities (one-step probability model), the individually-derived SR probabilities (SR model), as well as the combination of both of these factors (SR + one-step model; for details, see Methods). Figure 5c shows the results for the visual cortex ROI, plotting the relative AIC scores of all models across the eight TRs of interval trials (AIC_baseline_ -AIC_model_; negative values indicate better model fit compared to the baseline model). AIC scores indicated two phases of replay during the interval. In an early phase, beginning with the second TR after interval onset, we observed strongest evidence for reactivation of only the category that is one step away from the last shown stimulus (Fig. 5c, AICs at TR 2 for stimulus model *−*10709.42, 1-step model *−*10716.23, SR (multi-step) *−*10714.81, SR + 1-step *−*10715.90). In a later phase, between TR 4 to 6, and most strongly for TR 5, the model that jointly considered multi-step SR and one-step probabilities provided the best fit of classifier probabilities (Fig. 5c, AICs at TR 5 for stimulus model *−*15005.26, 1-step model *−*15007.24, SR (multi-step) *−*15005.54, SR + 1-step *−*15015.08). Hence, our results indicate on-task sequential reactivation of categories that were close to the last shown stimulus in participants’ mental maps of the task. To get a better understanding of the nature of this sequential reactivation, we extracted the residual classifier probabilities from the baseline model, i.e., the pattern of classifier probabilities that were above or below stimulus-driven activation resulting from the pre-interval trial history. Confirming our model fitting results, these residuals showed clear backwards ordering (Fig. 5d). Testing this ordering with the regression approach developed in Wittkuhn and Schuck (2021) showed a significant regression slope of sequential ordering at TRs 1–3 and 7 (Fig. 5e, *t*s *≥* 4.93, *p*s *< .*05, *d*s *≥* 0.79, *p*-values uncorrected). Our previous work has demonstrated that the time course of the regression slope indicates the direction and speed of neural replay (Wittkuhn and Schuck, 2021). In light of these findings, the observed change from a negative slope in the early TRs 1—3 to a reversed pattern by TR 7 is consistent with backwards replay occurring at a speed of 128 ms per item or slower.

We confirmed the above results by running an additional analysis that did not rely on modeling stimulus-driven influences, but rather used subsets of “clean” trials to assess the magnitude of reactivation in the absence of stimulus-driven influences. Leveraging the probabilistic nature of our task, we built a subset of the data in which all classifier probabilities were removed that reflected categories shown in the pre-trial history of the interval trial under consideration (minimum time since a category was last shown on the screen: 10 trials; time window sub-selection was done separately for each time point, participant and stimulus category based to our modeling approach, see Methods). This procedure reduced the number of trials available for further analysis to about 30% (19 out of 60 trials per participant) in the first TR and 80% (49 out of 60 trials) in the last TR of the on-task interval in the uni-directional graph condition (similar for the bi-directional graph condition; Fig. S11a). Applying the same regression approach used above, we again found a similar time course of sequentiality, with a significant regression slope of ordering at TRs 7–8 (Fig. 5f–g, *t*s *≥* 3.08, *p*s *< .*03, *d*s *≥* 0.49, *p*-values

FDR-corrected). As before, the ordering was indicative of fast backward replay as reflected in higher classifier probabilities for categories which lay in the past in the early TRs (TRs 1–3) followed by the reverse in the later interval TRs (TR 7). Repeating the same analyses in a motor cortex ROI revealed no comparable evidence for on-task replay in this region (see Fig. S12).

Having established the existence of on-task replay in visual cortex, but not motor cortex, we next asked how replay changed across learning (recall that every participant experienced both graph structures and the graph structure changed without prior announcement halfway through the task, see Fig. 1e). Previous theoretical accounts have suggested that replay might be particularly beneficial after the transition structure of the environment changes in order to update previous task representations (Wittkuhn et al., 2021). In order to investigate how replay changed across learning and whether it would be influenced by the change in graph structure, we split each block data into half and ran the same modeling approach as described above on the partitioned datasets and compared the AIC scores of the SR + 1-step model to the stimulus model. This analysis revealed a better fit of the SR + 1-step model compared to the stimulus model in the second half of the second run up to the second half of the third run (Fig. 6a).

**Figure 6:**
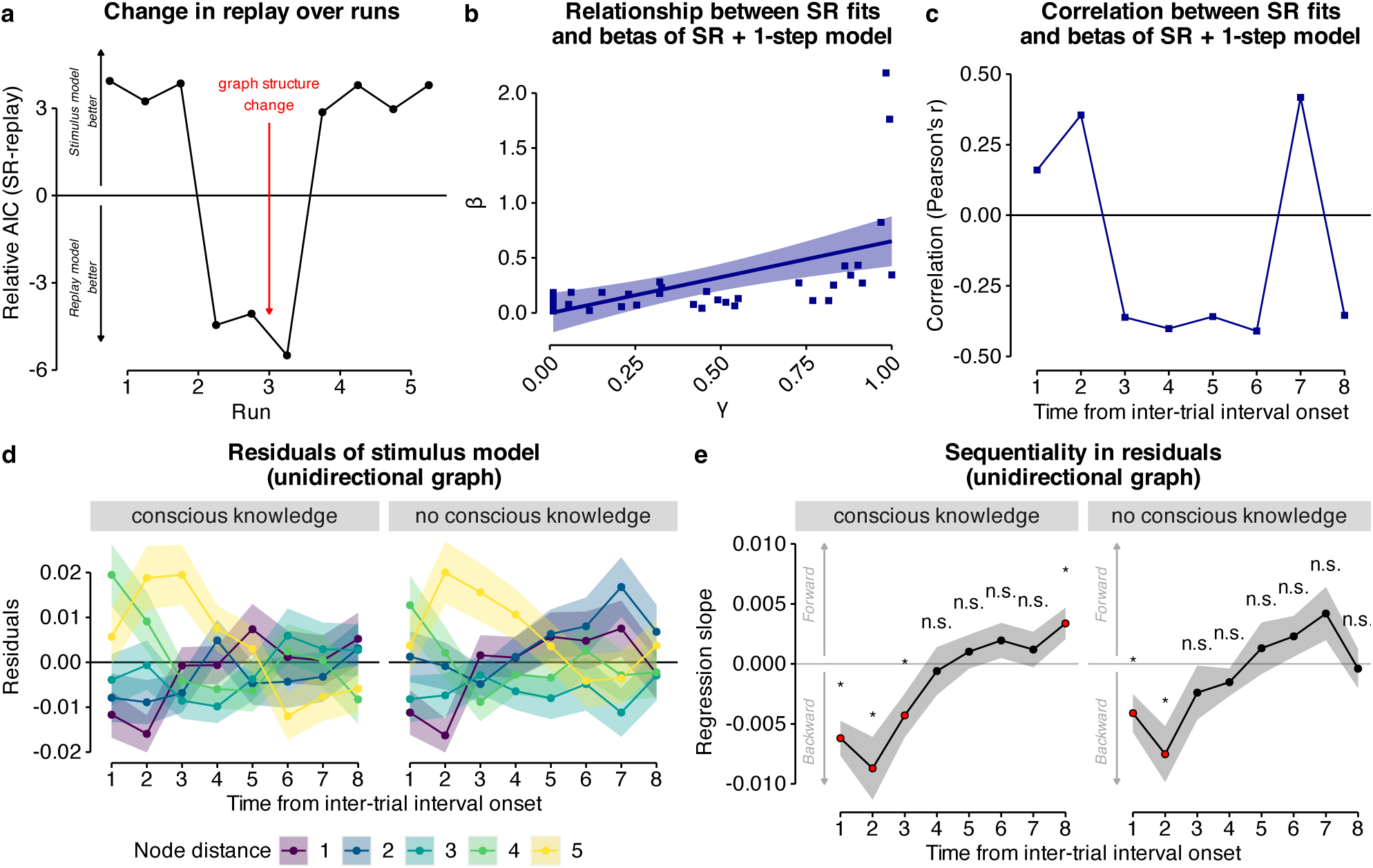
Time course of SR on-task replay, relationship between SR on-task replay and predictive depth of SR, and influence of conscious knowledge. **(a)** Time courses (runs; x-axis) of relative AIC scores (y-axis) comparing the full SR replay model against the baseline model in data from both uni- and bi-directional task conditions in the visual cortex ROI. The red arrow indicates the change in graph structure halfway through the third task run. Positive and negative values indicate better fit of the stimulus model and the SR replay model, respectively. **(b)** Relationship between the participant-wise behavioral model fit of the SR model (*γ* parameter; x-axis) and the participant-wise absolute betas of the regressors of the SR + 1-step model averaged across all eight TRs (y-axis). **(c)** Time courses (in TRs from inter-trial interval (ITI) onset; x-axis) of correlation coefficients (Pearson’s *r*; y-axis) quantifying the relationship between participant-wise betas of the regressors of the SR + 1-step model at each TR and the participant-wise behavioral model fit of the SR model (*γ* parameter) in data from the unidirectional task condition and visual cortex ROI. 1 TR = 1.25 s. All statistics have been derived from data of *n* = 39 human participants who participated in one experiment. **(d)** Time courses (in TRs from ITI onset; x-axis) of the residuals of each classifier node distance of the stimulus model (colors), split by self-reported conscious knowledge (plot as in Fig. 5d, unidirectional graph data). **(e)** Time courses (in TRs from ITI onset; x-axis) of mean regression slopes (y-axis) relating node distance (i.e., sequential position from current node in the graph structure) to their residuals in the stimulus model as in (d), split by self-reported conscious knowledge (“conscious knowledge” vs. “no conscious knowledge”; also see Fig. 2f). Positive and negative values indicate forward and backward sequentiality, respectively. Red dots and asterisks indicate significant differences from baseline (horizontal gray line at zero; all *p*s *≤ .*05, uncorrected; two-sided one-sample t-tests, one test per TR). Shaded areas in (d) and (e) represent *±*1 SEM. All statistics have been derived from data of *n* = 39 human participants who participated in one experiment. 1 TR = 1.25 s.

Next, we asked which relation neural on-task replay had to behavior, based on the idea that SRs can be updated through replay, rather than through online experience alone (Russek et al., 2017, 2021). To this end, we correlated averaged absolute beta values (across the entire interval trial) of the SR regressor from the SR + 1-step model in visual cortex with the participant-wise *γ* parameter of the behavioral SR model. This indicated a correlation of 0.55 and 0.46 for uni- and bi-directional graph structures, respectively (*p*s *< .*001, Fig. 6n). The correlation remained significant after removing outliers evident in Fig. 6b for data from the unidirectional graph (*r* = 0.64, *p < .*001; removing two participants with a large mean beta), but not the bidirectional graph (*r* = 0.25, *p* = .15; removing three participants with a large mean beta). We did not find a correlation between the *γ* parameter and the stimulus or 1-step regressors (*p ≤* 0.52 and *p ≤* 0.15, respectively; *p*-values uncorrected). A time-resolved analysis of the relationship between on-task SR replay and the behavioral SR parameter that correlated the individual fMRI SR regressor with the individually fitted behavioral SR *γ* parameter showed significant negative associations in TRs 3–6 and 8 (*r*s *≤ −.*35, *p*s *≤ .*05, FDR-corrected) and a significant positive correlation in TRs 2 and 7 (*r*s *≥ .*35, *p*s *< .*05, FDR-corrected) in unidirectional data in the visual cortex ROI (Fig. 6c).

Finally, we examined whether conscious task knowledge had an effect on on-task SR replay. To this end, we split the data of the stimulus model residuals shown in Fig. 5d by whether participants had reported conscious knowledge of the sequential task structure or not (“yes” vs. “no” response; see Fig. 2f). This analysis revealed no qualitative differences in the time course of SR replay between the two groups. Both groups showed clear backward ordering in early TRs, i.e., stronger activation of items likely to have preceded the shown stimulus (yellow line, Fig. 6d, compared to Fig. 5d), followed by the reverse pattern in late TRs. Using the same quantification of sequenceness as before (Fig. 5e), confirmed that both groups exhibited the same sequentiality dynamics, with significant negative (backwards) effects in early TRs in both groups, and positive effects in late TRs (significant only for the conscious group at TR 8, although the numeric values were very close, with *p_unc._* = .07 at TR 7 in the group without conscious awareness, see Fig. 6e). This suggests that on-task backward SR replay was unrelated to conscious knowledge.

## Discussion

In the current study, human participants performed an incidental statistical learning task. While only half of our sample reported conscious knowledge of the experienced sequential regularities, we found clear behavioral evidence that the majority of participants learned multi-step sequential expectations as predicted by a successor representation (SR) model. Our neuroimaging results show on-task backward replay in visual cortical areas while participants paused only briefly for 10 seconds during the ongoing task. Sequential replay was consistent with the SR model and correlated with behavioral evidence of SR model use, but independent of conscious knowledge.

Our fMRI sequentiality results are in line with our previous work (Schuck and Niv, 2019; Wittkuhn and Schuck, 2021), further validate our analytic approach, but also substantially extend our knowledge about replay in the human brain. The observed time course of the sequentiality slope suggested that on-task replay was backwards and, based on a comparison with data from Wittkuhn and Schuck (2021), occurred on a time scale on the order of 32 – 128 ms between item activations. The direction and speed of replay are notable given that they run counter to ideas that stipulate backwards replay is related to reward processing, while planing would engage forward replay Foster and Wilson (2006), which – in humans – could correspond to a slow conscious process of step-by-step thinking. In this light, we highlight three aspects of the replay observed here that differentiate our study from previous work: First, replay occurred during very short pauses from ongoing behavior, that differ substantially from the minute- or hour-long rest and sleep periods during which most replay has been detected and investigated in humans so far. Participants were also not instructed to learn about any sequentiality in the task and were not informed about the purpose of on task pauses, which furthermore occurred at unpredictable times. Second, we provide a test of whether replay is related to consciousness in humans and find no evidence for this idea. Third, replay occurred only in visual cortex, but not in motor cortex, even though the task required motor responses.

Our behavioral results are consistent with previous findings showing that humans learn about networks of stimuli beyond one-step transitions (e.g., Schapiro et al., 2013; Karuza et al., 2016, 2017, 2019; Garvert et al., 2017; Kahn et al., 2018; Lynn and Bassett, 2020; Lynn et al., 2020a,b). Computational modeling showed that an SR model with a medium predictive horizon best explained behavioral data, thereby establishing a link between previously known behavioral effects and online TD learning of an SR model (Dayan, 1993). Our findings add to a growing set of studies that uses models based on SRs to demonstrate the formation of predictive representations of task structure in human behavioral and neuroimaging data (Garvert et al., 2017; Russek et al., 2017; Momennejad et al., 2017; Momennejad, 2020; Russek et al., 2021; Garvert et al., 2023). We note that statistical regularities in our main task were governed by two graph structures: one for transitions in the first half of the experiment and another for the second half. Our results therefore speak to the idea that SR learning can be considered a continuous process that adapts to environmental changes. In a supplementary analysis we considered behavior separately for each graph condition and order, and found that the predictive horizon was influenced by learning history. During learning of the first graph structure the predictive horizon was at *γ* = 0.55, regardless of graph condition (uni or bi). After the transition structure changed to the second graph structure halfway through the task, however, the predictive horizon dependent on the order: participants who first learned the unidirectional and then the bidirectional graph, had a higher discount parameter of *γ* = 0.75, compared to participants who experienced the reverse order and had a lower discount parameter of *γ* = 0.3 (Fig. S5). This suggests that SR learning is not only ongoing, but that it’s depth also adapts to environmental structure. This idea relates to recent work suggesting that the brain may host SRs at varying predictive horizons in parallel (Momennejad and Howard, 2018; Brunec and Momennejad, 2021).

Our evidence for on-task replay relates to research in rodents, where time-compressed sequential place cell activations, called theta sequences, occur during active behavior (Foster and Wilson, 2007) and reflect multiple potential future trajectories when the animal pauses at a decision point (Johnson and Redish, 2007), or cycle between future trajectories during movement (Kay et al., 2020) possibly reflecting an online planning process. However, in contrast to previous studies in rodents and humans (Kaplan et al., 2020; Kurth-Nelson et al., 2016; Eldar et al., 2020), participants in our experiment likely did not engage in any explicit planning process, as discussed above.

One important aspect of our work is that we focused on cortical replay of predictive representations in visual (occipito-temporal) and sensorimotor (pre- and postcentral gyri) cortex. Previous work has largely focused on the hippocampus as a site of replay and as a potential brain region to host predictive cognitive maps (Garvert et al., 2017; Stachenfeld et al., 2017), while other studies have also emphasized the role of the prefrontal cortex (PFC) (Wilson et al., 2014; Schuck et al., 2016; Badre and Nee, 2018). Several fMRI studies demonstrated that hippocampal activity is modulated by stimulus predictability in sequential learning tasks (Strange et al., 2005; Harrison et al., 2006; Bornstein and Daw, 2012) and is related to the reinstatement of cortical task representations in visual cortex (Bosch et al., 2014; Hindy et al., 2016; Kok and Turk-Browne, 2018). Replay is known to occur throughout the brain (see e.g., Foster, 2017) but the functions of distributed replay events still remain to be further illuminated. Our findings shed light on the distribution of predictive representations and replay in the human brain, and suggest a specific involvement of sensory but not motor areas. Yet, which roles the hippocampus and PFC play in this process remains an open question.

Concerning the observed backward order, we note that previous research has found awake replay in both forward and backward order in rodents (Foster and Wilson, 2006; Diba and Buzsáki, 2007; Gupta et al., 2010) as well as in humans (Liu et al., 2021), and suggested that the directionality of replay may be tied to different functions, such as memory consolidation vs. value learning (e.g., Foster and Wilson, 2006; Ólafsdóttir et al., 2018; Liu et al., 2019; Wittkuhn et al., 2021). Neural sequences that have been associated with a prospective planning function are typically in forward order relative to the experienced sequence (Johnson and Redish, 2007; van der Meer and Redish, 2009; Pfeiffer and Foster, 2013; Wikenheiser and Redish, 2015b). However, as others have pointed out before (Kurth-Nelson et al., 2016), it is plausible to plan backward instead of forward (also see LaValle, 2006), and previous studies also reported backward sequences during theta in rodents (Wang et al., 2020) as well as during value learning in humans (Liu et al., 2021).

In conclusion, our results suggest a role of cortical replay in human unconscious learning and use of predictive maps of the environment during short on-task pauses.

## Methods

### Participants

44 young and healthy adults were recruited from an internal participant database or through local advertisement and fully completed the experiment. No statistical methods were used to predetermine the sample size but it was chosen to be larger than similar previous neuroimaging studies (e.g., Schuck and Niv, 2019; Momennejad et al., 2018; Tambini and Davachi, 2013). Five participants were excluded from further analysis because they viewed different task stimuli in session 1 and 2 due to a programming error in the behavioral task. Thus, the final sample consisted of 39 participants (mean age = 24.28 years, *SD* = 4.24 years, age range: 18 - 33 years, 23 female, 16 male). All participants were screened for MRI eligibility during a telephone screening prior to participation and again at the beginning of each study session according to standard MRI safety guidelines (e.g., asking for metal implants, claustrophobia, etc.). None of the participants reported to have any major physical or mental health problems. All participants were required to be right-handed, to have corrected-to-normal vision, and to speak German fluently. The ethics commission of the German Psychological Society (DGPs) approved the study protocol (reference number: SchuckNicolas2020-06-22VA). All volunteers gave written informed consent prior to the beginning of the experiments. Every participant received 70.00 Euro and a performance-based bonus of up to 5.00 Euro upon completion of the study. None of the participants reported to have any prior experience with the stimuli or the behavioral task.

### Task

#### Stimuli

All visual stimuli were taken from a set of colored and shaded images commissioned by Rossion and Pourtois (2004), which are loosely based on images from the original Snodgrass and Vanderwart set (Snodgrass and Vanderwart, 1980). The images are freely available on the internet at https://sites.google.com/andrew.cmu.edu/tarrlab/stimuli under the terms of the Creative Commons Attribution-NonCommercial-ShareAlike 3.0 Unported license (for details, see https://creativecommons.org/licenses/by-nc-sa/3.0/) and have been used in similar previous studies (e.g., Garvert et al., 2017). Stimulus images courtesy of Michael J. Tarr at Carnegie Mellon University (for details, see http://www.tarrlab.org/). In total, we selected 24 images which depicted animals that could be expected in a public zoo. Specifically, the images depicted a bear, a dromedary, a deer, an eagle, an elephant, a fox, a giraffe, a goat, a gorilla, a kangaroo, a leopard, a lion, an ostrich, an owl, a peacock, a penguin, a raccoon, a rhinoceros, a seal, a skunk, a swan, a tiger, a turtle, and a zebra (in alphabetical order; for an overview of all images, see Fig. S2). For each participant, six task stimuli were randomly selected from the set of the 24 animal images and each image was randomly assigned to one of six response buttons. This randomization ensured that any potential systematic differences between the stimuli (e.g., familiarity, preference, or ability to decode) would not influence the results on a group level (for a similar reasoning, see e.g., Liu et al., 2021). Cages were represented by a clipart illustration of a black fence which is freely available from https://commons.wikimedia.org/wiki/File:Maki-fence-15.svg, open-source and licensed under the Creative Commons CC0 1.0 Universal Public Domain Dedication, allowing further modification (for details, see https://creativecommons.org/publicdomain/zero/1.0/). When feedback was presented in the training and single trial task conditions, correct responses were indicated by a fence colored in green and incorrect responses were signaled by a fence colored in red. The color of the original image was modified accordingly. All stimuli were presented against a white background.

#### Hardware and software

Behavioral responses were collected using two 4-button inline fiber optic response pads (Current Designs, Philadelphia, PA, USA), one for each hand, with a linear arrangement of four buttons. Buttons were colored in blue, yellow, green, and red, from left to right, but participants were instructed that the button color was irrelevant for the task. For an illustration of the hand placement and response button mapping, see Fig. S3b. The two response pads were attached horizontally to a rectangular cushion that was placed in participants’ laps such that they could place their fingers on the response buttons with arms comfortably extended while resting on the scanner bed. Participants were asked to place their index, middle, and ring finger of their left and right hand on the yellow, green, and red buttons of the left and right response pads, respectively. The fourth (blue) button on each response pad was masked with tape and participants were instructed to never use this response button. Behavioral responses on the response pads were transferred to the computer running the experimental task and mapped to the keyboard keys z, g, r and w, n, d for the left and right hand, respectively. The task was programmed in PsychoPy3 (version 3.0.11; Peirce, 2007, 2008; Peirce et al., 2019) and run on a Windows 7 computer with a monitor refresh-rate of 16.7 ms. We recorded the presentation time stamps of all task events (onsets of all presentations of the fixation, stimulus, SRI, response, feedback, and ITI events) and confirmed that all components of the experimental task procedure were presented as expected.

#### Instructions

After participants entered the MRI scanner during the first study session and completed an anatomical T1-weighted (T1w) scan and a 5 min fMRI resting-state scan, they read the task instructions while lying inside the MRI scanner (for an illustration of the study procedure, see Fig. S1). Participants were asked to read all task instructions carefully (for the verbatim instructions, see Boxes S1 to S15). They were further instructed to clarify any potential questions with the study instructor right away and to lie as still and relaxed as possible for the entire duration of the MRI scanning procedure. As part of the instructions, participants were presented with a cover story in order to increase motivation and engagement (see Box S1). Participants were told to see themselves in the role of a zookeeper in training whose main task is to ensure that all animals are in the correct cages. In all task conditions, participants were asked to always keep their fingers on the response buttons to be able to respond as quickly and as accurately as possible. The full task instructions can be found in the supplementary information (SI), translated to English (see SI, starting on page 13, Boxes S1 to S15) from the original in German (see SI, starting at page 17).

#### Training trials

After participants read the instructions and clarified all remaining questions with the study instructors via the MRI intercom, they completed the *training* phase of the task (for an illustration of the trial procedure, see Fig. S3a). The training condition was designed to explicitly teach participants the assignment of stimuli to response buttons. Each of the six animal stimuli selected per participant was randomly assigned to one of six response buttons. For the training condition, participants were told to see themselves in the role of a zookeeper in training in a public zoo whose task is to learn which animal belongs in which cage (see Box S1). During each trial, participants saw six black cages at the bottom of the screen with each cage belonging to one of the six animals. On each trial, an animal appeared above one of the six cages. Participants were tasked to press the response button for that cage as fast and accurately as possible and actively remember the cage where the animal belonged (see Box S3 and Box S4). The task instructions emphasized that it would be very important for participants to actively remember which animal belonged in which cage and that they would have the chance to earn a higher bonus if they learned the assignment and responded accurately (see Box S5).

In total, participants completed 30 trials of the training condition. Across all trials, the pairwise ordering of stimuli was set to be balanced, with each pairwise sequential combination of stimuli presented exactly once, i.e., with *n* = 6 stimuli, this resulted in *n ∗* (*n −* 1) = 6 *∗* (6 *−* 1) = 30 trials. In this sense, the stimulus order was drawn from a random walk along the graph with all nodes connected to each other and an equal probability of *p_ij_* = 0.2 of transitioning from one node to any other node in the graph. This pairwise balancing of sequential combinations was used to ensure that participants would not learn any particular sequential order among the stimuli. Note, that this procedure only controlled for sequential order between pairs of consecutive stimuli but not higher-order sequential ordering of two steps or more.

On the first trial of the training condition, participants first saw a small black fixation cross that was displayed centrally on the screen for a fixed duration of 300 ms and signaled the onset of the following stimulus. The fixation cross was only shown on the first trial of the training phase, to allow for a short preparation signal before stimulus presentation began. Following the fixation cross, one of the animals was presented in the upper half of the screen above one of six cages that referred to the six response buttons and were presented in the lower half of the screen. The stimuli were shown for a fixed duration of 800 ms which was also the maximum time allowed for participants to respond. Note, that the instructions told participants that they would have 1 s to respond (see Box S4), an actual difference of 200 ms that was likely hardly noticeable by participants. Following the stimulus, participants always received feedback that was shown for a fixed duration of 500 ms. If participants responded correctly, the cage corresponding to the correctly pressed response button, was shown in green. If participants did not respond correctly, the cage referring to the correct response button was shown in green and the cage referring to the incorrectly pressed response button was shown in red. If participants responded too late, the cage referring to the correct response button was shown in green and the German words “Zu langsam” (in English: “Too slow”) appeared in large red letters in the upper half of the screen. Finally, a small black fixation cross was shown during an ITI with a variable duration of *M* = 1500 ms. The ITIs were drawn from a truncated exponential distribution with a mean of *M* = 1.5 s, a lower bound of *x*_1_ = 1.0 s and an upper bound of *x*_2_ = 10.0 s. To this end, we used the truncexpon distribution from the SciPy package (Virtanen et al., 2020) implemented in Python 3 (Van Rossum and Drake, 2009). The truncexpon distribution is described by three parameters, the shape *b*, the location *µ* and the scale *β*. The support of the distribution is defined by the lower and upper bounds, [*x*_1_*, x*_2_], where *x*_1_ = *µ* and *x*_2_ = *b ∗ β* + *µ*. We solved the latter equation for the shape *b* to get *b* = (*x*_2_ *− x*_1_)*/β*. We chose the scale parameter *β* such that the mean of the distribution would be *M* = 2.5. To this end, we applied scipy.optimize.fsolve (Virtanen et al., 2020) to a function of the scale *β* that becomes zero when *truncexpon.mean*((*x*_2_ *− x*_1_)*/β, µ, β*) *− M*) = 2.5. In total, the training phase took approximately 2 min to complete.

#### Single trials

After participants finished the training phase of the task in the first experimental session, they completed eight runs of the *single* condition and another ninth run at the beginning of the second session (for an illustration of the trial and study procedure, see Fig. 1a and Fig. S1, respectively). The single trial condition of the task mainly served two purposes: First, on a behavioral level, the single trial condition was used to further train participants on the associations between animal stimuli and response keys. Second, on a neural level, the single trial condition was designed to elicit object-specific neural activation patterns of the presented visual animal stimuli and the following motor response. The resulting neural activation patterns were later used to train the probabilistic classifiers (for details, see below). The cover story of the instructions told participants that they would be tested on how well they have learned the association between animals and response keys during the training phase (see Box S6).

In total, participants completed nine runs of the single trial condition. Eight runs were completed during session 1 and an additional ninth run was completed at the beginning of session 2 in order to remind participants about the S-R mappings (for an illustration of the study procedure, see Fig. S1). Each run consisted of 60 trials. As in the training phase, the proportion of pairwise sequential combinations of stimuli was balanced within a run. Across all trials, each pairwise sequential combination of stimuli was presented twice, i.e., with *n* = 6 stimuli, this results in *n ∗* (*n −* 1) *∗* 2 = 6 *∗* (6 *−* 1) *∗* 2 = 60 trials. As for the training trials, the sequential ordering of stimuli was drawn from a graph with all nodes connected to each other and an equal probability of *p_ij_*= 0.2 of transitioning from one node to any other node in the graph. With 60 trials per run, each of the six animal stimuli was shown 10 times per run. Given nine runs of the single trial condition in total, this amounted to a maximum of 90 trials per stimulus per participant of training examples for the classifiers. Including a ninth run at the beginning of session 2 offered two advantages. First, participants were reminded about the associations between the stimuli and response keys that they had learned extensively during session 1. Second, the ninth run allowed to investigate decoding performance across session boundaries. Note, that the two experimental sessions were separated by about one week. Although the pre-processing of fMRI data (for details, see section on fMRI data pre-processing below) should align the data of the two sessions, remaining differences between the two sessions (e.g., positioning of the participant in the MRI scanner) could lead to a decrement in decoding accuracy when testing classifiers that were trained on session 1 data to data from session 2. Our decoding approach was designed such that pattern classifiers would be mainly trained on neural data from single trials in session 1 but then applied to data from sequence trials in session 2.

As in training trials, the first trial of each run in the single trial phase started with a black fixation cross on a white background that was presented for a fixed duration of 300 ms. Only the first trial of a run contained a fixation cross, to provide a preparatory signal for participants which would later be substituted for by the ITI. Participants were then presented with one of the six animal stimuli that was presented centrally on the screen for a fixed duration of 500 ms. Participants were instructed to not respond to the stimulus (see instructions in Box S7). To check if participants indeed did not respond during the stimulus or the following SRI, we also recorded responses during these trial events. During the breaks between task runs, participants received feedback about the proportion of trials on which they responded too early. If participants responded too early, they were reminded by the study instructors to not respond before the response screen. A variable SRI followed the stimulus presentation during which a fixation cross was presented again. Including a jittered SRI ensured that the neural responses to the visual stimulus and the motor response could be separated in time and reduced temporal autocorrelation. Following the SRI, the cages indicating the response buttons were displayed centrally on the screen for a fixed duration of 800 ms, which was also the response time limit for participants. If participants responded incorrectly, the cage referring to the correct response button was shown in green and the cage referring to the incorrectly pressed response key was shown in red. If participants responded too late, the cage referring to the correct response button was shown in green and the German words “Zu langsam” (in English: “Too slow”) appeared in large red letters in the upper half of the screen. If participants responded correctly, the feedback screen was skipped. Each trial ended with an ITI with a variable duration of *M* = 2.5 s. Both SRIs and ITIs were drawn from a truncated exponential distribution as on training trials (for details, see the description of training trials above).

#### Sequence trials

Following the ninth run of the single trial condition in session 2, participants completed five runs of the *sequence* condition (for an illustration of the study procedure, see Fig. S1). During sequence trials, participants were exposed to a fast-paced stream of the same six animal stimuli as in the training and single trial phase. Unbeknownst to participants, the sequential ordering of animal stimuli now followed particular transition probabilities.

During the graph task, the sequential order of stimuli across trials was determined by two graph structures with distinct transition probabilities. In the first graph structure, each node had a high probability (*p_ij_* = 0.7) of transitioning to the next neighboring (i.e., transitioning from *A* to *B*, *B* to *C*, *C* to *D*, *D* to *E*, *E* to *F*, and *F* to *A*). Transitions to all other nodes (except the previous node) happened with equal probability of 0.1. Transitions to the previous node never occurred (transition probability of *p_ij_* = 0.0). These transition probabilities resulted in a sequential ordering of stimuli that can be characterized by a continuous progression in a unidirectional (i.e., clockwise) order around the ring-like graph structure. We therefore termed this graph structure the *unidirectional graph* (or *uni*, in short). The second graph structure allowed sequential ordering that could also progress in counterclockwise order. To this end, stimuli were now equally likely to transition to the next neighboring but also the previous node (probability of *p_ij_* = 0.35, i.e., splitting up the probability of *p_ij_* = 0.7 of transitioning to the next neighboring node only in the unidirectional graph structure). As in the unidirectional graph, transitions to all other nodes happened with equal probability of *p_ij_*= 0.1. Given that stimuli could follow a sequential ordering in both directions of the ring, we refer to this graph structure as the *bidirectional graph* (or *bi*, in short).

Participants completed five runs of the sequence trials. Each run consisted of 240 trials. Each stimulus was shown 40 times per run. In the unidirectional graph, for each stimulus the most likely transitions (probability of *p_ij_*= 0.7) to the next neighboring node occurred 28 times per participant. Per stimulus and participant, 4 transitions to the other three possible nodes (low probability of *p_ij_* = 0.1) happened. No transitions to the previous node happened when stimulus transitions were drawn from a unidirectional graph structure. Together, this resulted in 28 + 4 *∗* 3 = 40 presentations per stimulus, run and participant. For the bidirectional graph structure, transitions to the next neighboring and the previous node occurred 14 times per stimulus and to all other nodes 4 times as for the unidirectional graph structure. Together, this resulted in 14 + 14 + 4 *∗* 3 = 40 presentations per stimulus, run and participant.

As for the other task conditions, only the first trial of the sequence trial phase started with the presentation of a small black fixation cross that was presented centrally on the screen for a fixed duration of 300 ms. Then, an animal stimulus was presented centrally on the screen for a fixed duration of 800 ms, which also constituted the time limit in which participants could respond with the correct response button. Participants did not receive feedback during sequence trials in order to avoid any influence of feedback on sequence learning. The stimulus was followed by an ITI with a mean duration of 750 ms. The ITI in the sequence trials was also drawn from a truncated exponential distribution with a mean of *M* = 750 ms, a lower bound of *x*_1_ = 500 ms and an upper bound of *x*_2_ = 5000 ms.

Importantly, during the sequence trials, we also included long inter-trial intervals (ITIs) of 10 s in order to investigate on-task replay. As stated above, participants completed 240 trials of the sequence trials per run. In each run, each stimulus was shown on a total of 40 trials. For each stimulus, every 10^th^ trial on average was selected to be followed by a long ITI of 10 s. This meant that in each of the five main task runs, 4 trials per stimulus were followed by a long ITI. In total, each participant experienced 24 long ITI trials per run and 120 long ITI trials across the entire experiment. The duration of 10 s (roughly corresponding to eight TRs at a TR of 1.25 s) was chosen based on our previous results showing that the large majority of sequential fMRI signals can be captured within this time period (cf. Wittkuhn and Schuck, 2021, their Fig. 3).

#### Post-task questionnaire

After participants left the scanner in session 2, they were asked to complete a computerized post-task questionnaire consisting of four parts. First, participants were asked to report their handedness by selecting from three alternative options, “left”, “right” or “both”, in a forced-choice format. Note, that participants were required to be right-handed to participate in the study, hence this question merely served to record the self-reported handedness in addition to the participant details acquired as part of the recruitment procedure and demographic questionnaire assessment. Second, participants were asked whether they noticed any sequential order among the animal stimuli during sequence trials and could respond either “yes” or “no” in a forced-choice format. Third, if participants indicated that they had noticed a sequential order of the stimuli (i.e., if they answered “yes” to the previous question), they were asked to indicate during which run of the sequence trials they had started to notice the ordering (selecting from run “1” to “5”). In case participants indicated that they did not notice a sequential ordering, they were asked to select “None” when asked about the run. Fourth, participants were presented with all sequential combinations of pairs of the animal stimuli and asked to indicate how likely animal A (on the left) was followed by animal B (on the right) during the sequence trial condition of the task. Participants were instructed to “follow their gut feeling” in case they were uncertain about the probability ratings. With *n* = 6 stimuli, this resulted in *n∗*(*n−*1) = 6*∗*(6*−*1) = 30 trials. Participants indicated their response using a horizontal slider on a continuous scale from 0% to 100%. We recorded participants probability rating and response time on each trial. There was no time limit for any of the assessments in the questionnaire. Participants took *M* = 5.49 min (*SD* = 2.38 min; range: 2.23 to 12.63 min) to complete the questionnaire. The computerized questionnaire was programmed in PsychoPy3 (version 3.0.11; Peirce, 2007, 2008; Peirce et al., 2019) and run on the same Windows 7 computer that was used for the experimental task.

#### Study procedure

All participants were screened for study and MRI eligibility during a telephone screening prior to participation. The study consisted of two experimental sessions. For an illustration of the study procedure, see Fig. S1. As data collection took place during the COVID-19 pandemic, upon arrival at the study center in both sessions, participants were first asked about any symptoms that could indicate an infection with the SARS-CoV-2 virus. The study instructors then measured participants’ body temperature which was required to not be higher than 37.5°*C*. Participants were asked to read and sign all the relevant study documents at home prior to their arrival at the study center.

##### Session 1

The first MRI session (Fig. S1a) started with a short localizer sequence of ca. 1 min during which participants were asked to rest calmly, close their eyes and move as little as possible. Once the localizer data was acquired, the study staff aligned the field of view (FOV) for the acquisition of the T1w sequence. The acquisition of the T1w sequence took about 4 min to complete. Using the anatomical precision of the T1w images, the study staff then aligned the FOV for the functional MRI sequences. Here, the lower edge of the FOV was first aligned to the visually identified anterior commissure - posterior commissure (AC-PC) line of the participant’s brain. The FOV was then manually titled by 20 degrees forwards relative to the rostro-caudal axis (positive tilt; for details see the section on “MRI data acquisition” on page 27). Shortly before the functional MRI sequences were acquired, we performed Advanced Shimming. During the shimming period, which took ca. 2 min, participants were again instructed to move as little as possible and additionally asked to avoid swallowing to further reduce any potential movements. Next, we acquired functional MRI data during a resting-state period of 5 min. For this phase, participants were instructed to keep their eyes open and fixate a white fixation cross that was presented on a black background. Acquiring fMRI resting-state data before participants had any exposure to the task (including instructions) allowed us to record a resting-state period that was guaranteed to be free of any task-related neural activation or reactivation. Following this pre-task resting-state scan, participants read the task instructions inside the MRI scanner and were able to clarify any questions with the study instructions via the intercom system. Participants then performed the training phase of the task (for details, see the section “Training trials” on page 21; Fig. S3) while undergoing acquisition of functional MRI data. The training phase took ca. 2 min to complete. Following the training phase, participants performed eight runs of the single trial phase of the task of ca. 6 min each (for details, see section “Single trials” on page 23; Fig. 1a) while fMRI data was recorded. Before participants left the scanner, field maps were acquired.

##### Session 2

At the beginning of the second session (Fig. S1b), participants first completed the questionnaire for MRI eligibility and the questionnaire on COVID-19 symptoms before entering the MRI scanner again. As in the first session, the second MRI session started with the acquisition of a short localizer sequence and a T1w sequence followed by the orientation of the FOV for the functional acquisitions and the Advanced Shimming. Participants were asked to rest calmly and keep their eyes closed during this period. Next, during the first functional sequence of the second study session, participants performed a ninth run of the single trial phase of the task in order to remind them about the correct response buttons associated with each of the six stimuli. We then acquired functional resting-state scans of 3 min each and functional task scans of 10 min each in an interleaved fashion, starting with a resting-state scan. During the acquisition of functional resting-state data, participants were asked to rest calmly and fixate a small white cross on a black background that was presented on the screen. During each of the functional task scans, participants performed the sequence task (for details, see section “Sequence trials” on page 24; Fig. 1b). Importantly, half-way through the third block of the sequence task, the graph structure was changed without prior announcement towards the second graph structure. After the sixth resting-state acquisition, field maps were acquired and participants eventually left the MRI scanner.

#### MRI data acquisition

All MRI data were acquired using a 32-channel head coil on a research-dedicated 3-Tesla Siemens Magnetom TrioTim MRI scanner (Siemens, Erlangen, Germany) located at the Max Planck Institute for Human Development in Berlin, Germany.

At the beginning of each of the two MRI recording sessions, high-resolution T1w anatomical Magnetization Prepared Rapid Gradient Echo (MPRAGE) sequences were obtained from each participant to allow co-registration and brain surface reconstruction (sequence specification: 256 slices; TR = 1900 ms; echo time (TE) = 2.52 ms; flip angle (FA) = 9 degrees; inversion time (TI) = 900 ms; matrix size = 192 x 256; FOV = 192 x 256 mm; voxel size = 1 x 1 x 1 mm).

For the functional scans, whole-brain images were acquired using a segmented k-space and steady state T2*-weighted multi-band (MB) echo-planar imaging (EPI) single-echo gradient sequence that is sensitive to the blood-oxygen-level dependent (BOLD) contrast. This measures local magnetic changes caused by changes in blood oxygenation that accompany neural activity (sequence specification: 64 slices in interleaved ascending order; anterior-to-posterior (A-P) phase encoding direction; TR = 1250 ms; TE = 26 ms; voxel size = 2 x 2 x 2 mm; matrix = 96 x 96; FOV = 192 x 192 mm; FA = 71 degrees; distance factor = 0%; MB acceleration factor 4). Slices were tilted for each participant by 20 degrees forwards relative to the rostro-caudal axis (positive tilt) to improve the quality of fMRI signal from the hippocampus (cf. Weiskopf et al., 2006) while preserving sufficient coverage of occipito-temporal and motor brain regions. The same sequence parameters were used for all acquisitions of fMRI data. For each functional task run, the task began after the acquisition of the first four volumes (i.e., after 5.00 s) to avoid partial saturation effects and allow for scanner equilibrium.

The first MRI session included nine functional task runs in total. After participants read the task instructions inside the MRI scanner, they completed the training trials of the task which explicitly taught participants the correct mapping between stimuli and response keys. During this task phase, 80 volumes of fMRI were collected, which were not used in any further analysis. The other eight functional task runs during session 1 consisted of eight runs of the single trial condition. Each run of the single trial task was about 6 min in length, during which 320 functional volumes were acquired. We also recorded two functional runs of resting-state fMRI data, one before and one after the task runs. Each resting-state run was about 5 min in length, during which 233 functional volumes were acquired.

The second MRI session included six functional task runs in total. After participants entered the MRI scanner, they completed a ninth run of the single trial task. As before, this run of the single trial task was also about 6 min in length, during which 320 functional volumes were acquired. Participants then completed five runs of the sequence learning task. Each run of the five sequence learning runs was about 10 min in length, during which 640 functional volumes were acquired. The five runs of the sequence learning task were interleaved with six recordings of resting-state fMRI data, each about 3 min in length, during which 137 functional volumes were acquired.

At the end of each scanning session, two short acquisitions with six volumes each were collected using the same sequence parameters as for the functional scans but with varying phase encoding polarities, resulting in pairs of images with distortions going in opposite directions between the two acquisitions (also known as the *blip-up / blip-down* technique). From these pairs the displacement maps were estimated and used to correct for geometric distortions due to susceptibility-induced field inhomogeneities as implemented in the fMRIPrep preprocessing pipeline (Esteban et al., 2018, for details, see below). In addition, a whole-brain spoiled gradient recalled (GR) field map with dual echo-time images (sequence specification: 36 slices; A-P phase encoding direction; TR = 400 ms; TE1 = 4.92 ms; TE2 = 7.38 ms; FA = 60 degrees; matrix size = 64 x 64; FOV = 192 x 192 mm; voxel size = 3 x 3 x 3.75 mm) was obtained as a potential alternative to the blip-up / blip-down method described above.

We also measured respiration during each scanning session using a pneumatic respiration belt as part of the Siemens Physiological Measurement Unit (PMU). Pulse data could not be recorded as the recording device could not be attached to the participants’ index finger as it would have otherwise interfered with the motor responses using the index finger (cf. Fig. S3b).

#### MRI data preparation

##### Arrangement of data according to the Brain Imaging Data Structure (BIDS) standard

The majority of the steps involved in preparing and preprocessing the MRI data employed recently developed tools and workflows aimed at enhancing standardization and reproducibility of task-based fMRI studies (for a similar data processing pipeline, see e.g., Esteban et al., 2019a; Wittkuhn and Schuck, 2021). Version-controlled data and code management was performed using DataLad (version 0.13.0; Halchenko et al., 2019, 2021), supported by the DataLad handbook (Wagner et al., 2020). Following successful acquisition, all study data were arranged according to the brain imaging data structure (BIDS) specification (Gorgolewski et al., 2016) using the HeuDiConv tool (version 0.8.0.2; freely available from https://github.com/ReproNim/reproin or https://hub.docker.com/r/repronim/ reproin) in combination with the ReproIn heuristic (Visconti di Oleggio Castello et al., 2020, version 0.6.0) that allows automated creation of brain imaging data structure (BIDS) data sets from the acquired Digital Imaging and Communications in Medicine (DICOM) images. To this end, the sequence protocol of the MRI data acquisition was set up to conform with the specification required by the ReproIn heuristic (for details of the heuristic, see https://github.com/nipy/heudiconv/ blob/master/heudiconv/heuristics/reproin.py). HeuDiConv was run inside a Singularity container (Kurtzer et al., 2017; Sochat et al., 2017) that was built from the most recent version (at the time of access) of a Docker container (tag 0.8.0.2), available from https://hub.docker.com/r/ repronim/reproin/tags. DICOMs were converted to the NIfTI-1 format using dcm2niix (version 1.0.20190410*GCC*6.3.0; Li et al., 2016). In order to make personal identification of study participants unlikely, we eliminated facial features from all high-resolution structural images using pydeface (version 2.0.0; Gulban et al., 2019, available from https://github.com/poldracklab/pydeface or https://hub.docker.com/r/poldracklab/pydeface). pydeface (Gulban et al., 2019) was run inside a Singularity container (Kurtzer et al., 2017; Sochat et al., 2017) that was built from the most recent version (at the time of access) of a Docker container (tag 37-2e0c2d), available from https://hub.docker.com/r/poldracklab/pydeface/tags and used Nipype, version 1.3.0-rc1 (Gorgolewski et al., 2011, 2019). During the process of converting the study data to BIDS the data set was queried using pybids (version 0.12.1; Yarkoni et al., 2019a,b), and validated using the bids-validator (version 1.5.4; Gorgolewski et al., 2020). The bids-validator (Gorgolewski et al., 2020) was run inside a Singularity container (Kurtzer et al., 2017; Sochat et al., 2017) that was built from the most recent version (at the time of access) of a Docker container (tag v1.5.4), available from https://hub.docker.com/r/bids/validator/tags.

##### MRI data quality control

The data quality of all functional and structural acquisitions was evaluated using the automated quality assessment tool MRIQC, version 0.15.2rc1 (for details, see Esteban et al., 2017, and the MRIQC documentation, available at https://mriqc.readthedocs.io/en/stable/). The visual group-level reports of the estimated image quality metrics confirmed that the overall MRI signal quality of both anatomical and functional scans was highly consistent across participants and runs within each participant.

##### MRI data preprocessing

Preprocessing of MRI data was performed using fMRIPrep 20.2.0 (long-term support (LTS) release; Esteban et al., 2018, 2019b, RRID:SCR 016216), which is based on Nipype 1.5.1 (Gorgolewski et al., 2011, 2019, RRID:SCR 002502). Many internal operations of fMRIPrep use Nilearn 0.6.2 (Abraham et al., 2014, RRID:SCR 001362), mostly within the functional processing workflow. For more details of the pipeline, see the section corresponding to workflows in fMRIPrep’s documentation at https://fmriprep.readthedocs.io/en/latest/workflows.html. Note, that version 20.2.0 of fMRIPrep is a long-term support (LTS) release, offering long-term support and maintenance for four years.

##### Preprocessing of anatomical MRI data using **fMRIPrep**

A total of two T1w images were found within the input BIDS data set, one from each study session. All of them were corrected for intensity non-uniformity (INU) using N4BiasFieldCorrection (Tustison et al., 2010), distributed with Advanced Normalization Tools (ANTs) 2.3.3 (Avants et al., 2008, RRID:SCR 004757). The T1w-reference was then skull-stripped with a Nipype implementation of the antsBrainExtraction.sh workflow (from ANTs), using OASIS30ANTs as target template. Brain tissue segmentation of cerebrospinal fluid (CSF), white-matter (WM) and gray-matter (GM) was performed on the brain-extracted T1w using fast (FMRIB Software Library (FSL) 5.0.9, RRID:SCR 002823, Zhang et al., 2001). A T1w-reference map was computed after registration of two T1w images (after INU-correction) using mri robust template (FreeSurfer 6.0.1, Reuter et al., 2010). Brain surfaces were reconstructed using recon-all (FreeSurfer 6.0.1, RRID:SCR 001847, Dale et al., 1999), and the brain mask estimated previously was refined with a custom variation of the method to reconcile ANTs-derived and FreeSurfer-derived segmentations of the cortical GM of Mindboggle (RRID:SCR 002438, Klein et al., 2017). Volume-based spatial normalization to two standard spaces (MNI152NLin6Asym, MNI152NLin2009cAsym) was performed through nonlinear registration with antsRegistration (ANTs 2.3.3), using brain-extracted versions of both T1w reference and the T1w template. The following templates were selected for spatial normalization: FSL’s MNI ICBM 152 non-linear 6^th^ Generation Asymmetric Average Brain Stereotaxic Registration Model (Evans et al., 2012, RRID:SCR 002823; TemplateFlow ID: MNI152NLin6Asym), ICBM 152 Nonlinear Asymmetrical template version 2009c (Fonov et al., 2009, RRID:SCR 008796; TemplateFlow ID: MNI152NLin2009cAsym).

##### Preprocessing of functional MRI data using **fMRIPrep**

For each of the BOLD runs found per participant (across all tasks and sessions), the following pre-processing was performed. First, a reference volume and its skull-stripped version were generated using a custom methodology of fMRIPrep. A B0-nonuniformity map (or fieldmap) was estimated based on two (or more) echo-planar imaging (EPI) references with opposing phase-encoding directions, with 3dQwarp (Cox and Hyde, 1997, AFNI 20160207). Based on the estimated susceptibility distortion, a corrected echo-planar imaging (EPI) reference was calculated for a more accurate coregistration with the anatomical reference. The BOLD reference was then co-registered to the T1w reference using bbregister (FreeSurfer) which implements boundary-based registration (Greve and Fischl, 2009). Co-registration was configured with six degrees of freedom. Head-motion parameters with respect to the BOLD reference (transformation matrices, and six corresponding rotation and translation parameters) are estimated before any spatiotemporal filtering using mcflirt (FSL 5.0.9, Jenkinson et al., 2002). BOLD runs were slice-time corrected using 3dTshift from AFNI 20160207 (Cox and Hyde, 1997, RRID:SCR 005927). The BOLD time-series were resampled onto the following surfaces (FreeSurfer reconstruction nomenclature): fsnative. The BOLD time-series (including slice-timing correction) were resampled onto their original, native space by applying a single, composite transform to correct for head-motion and susceptibility distortions. These resampled BOLD time-series will be referred to as preprocessed BOLD in original space, or just preprocessed BOLD. The BOLD time-series were resampled into standard space, generating a preprocessed BOLD run in MNI152NLin6Asym space. First, a reference volume and its skull-stripped version were generated using a custom methodology of fMRIPrep. Several confounding time-series were calculated based on the preprocessed BOLD: framewise displacement (FD), DVARS and three region-wise global signals. FD was computed using two formulations following Power et al. (absolute sum of relative motions, 2014) and Jenkinson et al. (relative root mean square displacement between affines, 2002). FD and DVARS are calculated for each functional run, both using their implementations in Nipype (following the definitions by Power et al., 2014). The three global signals are extracted within the CSF, the WM, and the whole-brain masks. Additionally, a set of physiological regressors were extracted to allow for component-based noise correction (CompCor, Behzadi et al., 2007). Principal components are estimated after high-pass filtering the preprocessed BOLD time-series (using a discrete cosine filter with 128*s* cut-off) for the two CompCor variants: temporal (tCompCor) and anatomical (aCompCor). tCompCor components are then calculated from the top 2% variable voxels within the brain mask. For aCompCor, three probabilistic masks (CSF, WM and combined CSF+WM) are generated in anatomical space. The implementation differs from that of Behzadi et al. (2007) in that instead of eroding the masks by 2 pixels on BOLD space, the aCompCor masks are subtracted from a mask of pixels that likely contain a volume fraction of GM. This mask is obtained by dilating a GM mask extracted from the FreeSurfer’s aseg segmentation, and it ensures components are not extracted from voxels containing a minimal fraction of GM. Finally, the masks are resampled into BOLD space and binarized by thresholding at 0.99 (as in the original implementation). Components are also calculated separately within the WM and CSF masks. For each CompCor decomposition, the *k* components with the largest singular values are retained, such that the retained components’ time series are sufficient to explain 50 percent of variance across the nuisance mask (CSF, WM, combined, or temporal). The remaining components are dropped from consideration. The head-motion estimates calculated in the correction step were also placed within the corresponding confounds file. The confound time series derived from head motion estimates and global signals were expanded with the inclusion of temporal derivatives and quadratic terms for each (Satterthwaite et al., 2013). Frames that exceeded a threshold of 0.5 mm FD or 1.5 standardized DVARS were annotated as motion outliers. All resamplings can be performed with a single interpolation step by composing all the pertinent transformations (i.e. headmotion transform matrices, susceptibility distortion correction when available, and co-registrations to anatomical and output spaces). Gridded (volumetric) resamplings were performed using antsApplyTransforms (ANTs), configured with Lanczos interpolation to minimize the smoothing effects of other kernels (Lanczos, 1964). Non-gridded (surface) resamplings were performed using mri vol2surf (FreeSurfer).

##### Additional preprocessing of functional MRI data following **fMRIPrep**

Following preprocessing using fMRIPrep, the fMRI data were spatially smoothed using a Gaussian mask with a standard deviation (Full Width at Half Maximum (FWHM) parameter) set to 4 mm using an example Nipype smoothing workflow ( for details, see the Nipype documentation) based on the Smallest Univalue Segment Assimilating Nucleus (SUSAN) algorithm as implemented in FSL (Smith and Brady, 1997). In this workflow, each run of fMRI data was separately smoothed using FSL’s SUSAN algorithm with the brightness threshold set to 75% of the median value of each run and a mask constituting the mean functional image of each run.

##### Multi-variate fMRI pattern analysis

All fMRI pattern classification analyses were conducted using the open-source Python (Python Software Foundation, Python Language Reference, version 3.8.6) packages Nilearn (version 0.7.0; Abraham et al., 2014) and scikit-learn (version 0.24.1; Pedregosa et al., 2011). In all classification analyses, we trained an ensemble of six independent classifiers, one for each of the six event classes. Depending on the analysis, these six classes either referred to the identity of the six visual animal stimuli or the identity of the participant’s motor response, when training the classifiers with respect to the stimulus or the motor onset, respectively. For each class-specific classifier, labels of all other classes in the data were relabeled to a common “other” category. In order to ensure that the classifier estimates were not biased by relative differences in class frequency in the training set, the weights associated with each class were adjusted inversely proportional to the class frequencies in each training fold. Given that there were six classes to decode, the frequencies used to adjust the classifiers’ weights were <u>^1^</u> for the class of interest, and <u>^5^</u> for the “other” class, comprising any other classes. Adjustments to minor imbalances caused by the exclusion of erroneous trials were performed in the same way. We used separate logistic regression classifiers with identical parameter settings. All classifiers were regularized using L2 regularization. The *C* parameter of the cost function was fixed at the default value of *C* = 1.0 for all participants. The classifiers employed the lbfgs algorithm to solve the multi-class optimization problem and were allowed to take a maximum of 4, 000 iterations to converge. Pattern classification was performed within each participant separately, never across participants. For each example in the training set, we added 4 s to the event onset and chose the volume closest to that time point (i.e., rounding to the nearest volume) to center the classifier training on the expected peaks of the BOLD response (i.e., accounting for hemodynamic lag; for a similar approach, see e.g., Deuker et al., 2013). At a TR of 1.25 s this corresponded roughly to the fourth MRI volume which thus compromised a time window of 3.75 s to 5.0 s after each event onset. We detrended the fMRI data separately for each run across all task conditions to remove low frequency signal intensity drifts in the data due to noise from the MRI scanner. For each classifier and run, the features were standardized (*z*-scored) by removing the mean and scaling to unit variance separately for each training and test set.

##### Classification procedures

First, in order to assess the ability of the classifiers to decode the correct class from fMRI patterns, we conducted a leave-one-run-out cross-validation procedure for which data from seven task runs of the single trials in session 1 were used for training and data from the left-out run (i.e., the eighth run) from session 1 was used for testing the classification performance. This procedure was repeated eight times so that each task run served as the testing set once. Classifier training was performed on data from all correct single trials of the seven runs in the respective cross-validation fold. Note that category order was randomized and trials were sufficiently separated to reduce temporal autocorrelation (SRIs and ITIs each 2500 ms, see procedure of single trials, cf. Dale, 1999; Wittkuhn and Schuck, 2021). In each iteration of the leave-one-run-out procedure, the classifiers trained on seven out of eight runs were then applied separately to the data from the left-out run. Specifically, the classifiers were applied to (1) data from the single trials of the left-out run, selecting volumes capturing the expected activation peaks to determine classification accuracy, and (2) data from the single trials of the left-out run, selecting all volumes from the volume closest to the stimulus or response onset and the next fifteen volumes to characterize temporal dynamics of probabilistic classifier predictions on a single trial basis. Second, we assessed decoding performance on single trials across the two experimental sessions. The large majority of fMRI data that was used to train the classifiers was collected in session 1 (eight of nine runs of the single trials), but the trained classifiers were mainly applied to fMRI data from session 2 (i.e., on-task intervals during sequence trials). At the beginning of the second experimental session, participants completed another run of the single trials (i.e., a ninth run; for the study procedure, see Fig. S1). This additional task run mainly served the two purposes of (1) reminding participants about the correct S-R mapping that they had learned in session 1, and (2) to investigate the ability of the classifiers to correctly decode fMRI patterns in session 2 when they were only trained on session 1 data. This second aspect is crucial, as the main focus of investigation is the potential reactivation of neural task representations in session 2 fMRI data. Thus, it is important to demonstrate that this ability is not influenced by losses in decoding performance due to decoding across session boundaries. In order to test cross-session decoding, we thus trained the classifiers on all eight runs of the single trial condition in session 1 and tested their decoding performance on the ninth run of the single trial condition in session 2. Classifiers trained on data from all nine runs of the single trials were subsequently applied to data from on-task intervals in sequence trials in session 2. For the classification analyses in on-task intervals of the sequence task, classifiers were trained on the peak activation patterns from all correct single trials (including session 1 and session 2 data) and then tested on all TR corresponding to the sequence task ITIs.

##### Feature selection

All participant-specific anatomical masks were created based on automated anatomical labeling of brain surface reconstructions from the individual T1w reference image created with Freesurfer’s reconall (Dale et al., 1999) as part of the fMRIPrep workflow (Esteban et al., 2018), in order to account for individual variability in macroscopic anatomy and to allow reliable labeling (Fischl et al., 2004; Poldrack, 2007). For the anatomical masks of occipito-temporal regions we selected the corresponding labels of the cuneus, lateral occipital sulcus, pericalcarine gyrus, superior parietal lobule, lingual gyrus, inferior parietal lobule, fusiform gyrus, inferior temporal gyrus, parahippocampal gyrus, and the middle temporal gyrus (cf. Haxby et al., 2001; Wittkuhn and Schuck, 2021). For the anatomical ROI of motor cortex, we selected the labels of the left and right gyrus precentralis as well as gyrus postcentralis. The labels of each ROI are listed in Table 1. Only gray-matter voxels were included in the generation of the masks as BOLD signal from non-gray-matter voxels cannot be generally interpreted as neural activity (Kunz et al., 2018). Note, however, that due to the whole-brain smoothing performed during preprocessing, voxel activation from brain regions outside the anatomical mask but within the sphere of the smoothing kernel might have entered the anatomical mask (thus, in principle, also including signal from surrounding non-gray-matter voxels).

**Table 1:** Labels used to index brain regions to create participant-specific anatomical masks of selected ROIs based on Freesurfer’s recon-all labels (Dale et al., 1999)

| ROI | Freesurfer labels (brain region) |
| --- | --- |
| Occipito-temporal | 1005, 2005 (cuneus); 1011, 2011 (lateral occipital sulcus); 1021, 2021 (pericalcarine gyrus); 1029, 2029 (superio parietal lobule); 1013, 2013 (lingual gyrus); 1008, 2008 (inferior parietal lobule); 1007, 2007 (fusiform gyrus); 1009, 2009 (inferior temporal gyrus); 1016, 2016 (parahippocampal gyrus); 1015, 2015 (middle temporal gyrus) |
| Motor | 1024, 2024 (left and right gyrus precentralis); 1022, 2022 (left and right gyrus postcentralis) |

### Statistical analyses

All statistical analyses were run inside a Docker software container or, if analyses were executed on a high performance computing (HPC), a Singularity version of the same container (Kurtzer et al., 2017; Sochat et al., 2017). All main statistical analyses were conducted using LME models employing the lmer function of the lme4 package (version 1.1.27.1, Bates et al., 2015) in R (version 4.1.2, R Core Team, 2019). If not stated otherwise, all models were fit with participants considered as a random effect on both the intercept and slopes of the fixed effects, in accordance with results from Barr et al. (2013) who recommend to fit the most complex model consistent with the experimental design. If applicable, explanatory variables were standardized to a mean of zero and a standard deviation of one before they entered the models. If necessary, we removed by-participant random slopes from the random effects structure to achieve a non-singular fit of the model (Barr et al., 2013). Models were fitted using the Bound Optimization BY Quadratic Approximation (BOBYQA) optimizer (Powell, 2007, 2009) with a maximum of 500, 000 function evaluations and no calculation of gradient and Hessian of nonlinear optimization solution. The likelihoods of the fitted models were assessed using Type III analysis of variance (ANOVA) with Satterthwaite’s method. A single-step multiple comparison procedure between the means of the relevant factor levels was conducted using Tukey’s honest significant difference (HSD) test (Tukey, 1949), as implemented in the emmeans package in R (version 1.7.0, Lenth, 2019; R Core Team, 2019). In all other analyses, we used one-sample *t*-tests if group data was compared to a baseline or paired *t*-tests if two samples from the same population were compared. If applicable, correction for multiple hypothesis testing was performed using the FDR (Benjamini and Hochberg, 1995) or Bonferroni (Bonferroni, 1936) correction method. If not stated otherwise, the *α*-level was set to *α* = 0.05, and analyses of response times included data from correct trials only. When effects of stimulus transitions during sequence trials were analyzed, data from the first trial of each run and the first trial after the change in transition structure in the third task run were removed.

### Statistical analyses of behavioral data

In order to test the a-priori hypothesis that behavioral accuracy in each of the nine runs of the single trials and five runs of the sequence trials would be higher than the chance level, we performed a series of one-sided one-sample *t*-tests that compared participants’ mean behavioral accuracy per run against the chance level of 100%*/*6 = 16.67% (Fig. S4). Participants’ behavioral accuracy was calculated as the proportion of correct responses per run (in %). The effect sizes (Cohen’s *d*) were calculated as the difference between the mean of behavioral accuracy scores across participants and the chance baseline (16.67%), divided by the standard deviation of the data (Cohen, 1988). The resulting *p*-values were adjusted for multiple comparisons using the Bonferroni correction (Bonferroni, 1936).

To examine the effect of task run on behavioral accuracy (Fig. S4b) and response times (Fig. 2b) in sequence trials, we conducted an LME model that included all five runs of sequence trials as a numeric predictor variable (runs 1 to 5) as the main fixed effect of interest as well as by-participant random intercepts and slopes.

Analyzing the effect of one-step transition probabilities on behavioral accuracy (Fig. 2c) and response times (Fig. 2d), we conducted two-sided paired *t*-tests comparing the effect of high vs. low transition probability separately for both unidirectional (*p_ij_* = 0.7 vs. *p_ij_* = 0.1) and bidirectional (*p_ij_* = 0.35 vs. *p_ij_* = 0.1) data. Effect sizes (Cohen’s *d*) were calculated by dividing the mean difference of the paired samples by the standard deviation of the difference (Cohen, 1988) and *p*-values were adjusted for multiple comparisons across both graph conditions and response variables using the Bonferroni correction (Bonferroni, 1936).

In order to examine the effect of node distance on response times in sequence trials (Fig. 2e), we conducted separate LME models for data from the unidirectional and bidirectional graph structures. For LME models of response time in unidirectional data, we included a linear predictor variable of node distance (assuming a linear increase of response time with node distance; see Fig. 1f, top right) as well as random intercepts and slopes for each participant. The linear predictor variable was coded such that the node distance linearly increased from *−*2 to +2 in steps of 1, modeling the hypothesized increase of response time with node distance from 1 to 5 (centered on the node distance of 3). For LME models of response time in bidirectional data, we included a quadratic predictor variable of node distance (assuming an inverted U-shaped relationship between node distance and response time; see Fig. 1f, bottom right) as well as by-participant random intercepts and slopes. The quadratic predictor variable of node distance was obtained by squaring the linear predictor variable.

To assess whether participants with and without conscious knowledge of sequential structure differed regarding the effect of node distance on response times, we conducted a series of two-sided two-sample *t*-tests comparing the effect of conscious knowledge (“yes” vs. “no”) separately for both unidirectional and bidirectional data. Effect sizes (Cohen’s *d*) were calculated by dividing the mean difference between the groups by the pooled standard deviation (Cohen, 1988) and *p*-values were adjusted for multiple comparisons across both graph conditions and response variables using the FDR correction (Benjamini and Hochberg, 1995).

### LME modeling of response times based on the successor representation

We modeled successor representations (SRs) for each participant depending on the transitions that were experienced in the task, including training, single and sequence trials. Specifically, each of the six stimuli was associated with a vector that reflected a *running* estimate of the long-term visitation probability of all six stimuli, starting from the current node. The successor matrix **M***^t^* was therefore a 6-by-6 matrix that contained six predictive vectors, one for each stimulus, and changed over time (hence the index *t*). The SR matrix on the first trial was initialized with a baseline expectation of 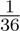 for each node. After a transition between stimuli *s_t_* and *s_t_*_+1_, the matrix row corresponding to *s_t_* was updated following a temporal difference (TD) learning rule (Dayan, 1993; Russek et al., 2017) as follows

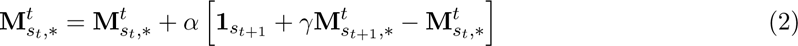

whereby **1***_st_*_+1_ is a one-hot vector with a 1 in the *s_t_*_+1_^th^ position, **M***^t^* is the row corresponding to stimulus *s_t_* of matrix 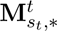. The learning rate *α* was arbitrarily set to a fixed value of *α* = 0.1, and the discount parameter *γ* was varied in increments of 0.05 from 0 to 0.95, as also described in the main text. This meant that the SR matrix would change throughout the task to reflect the experienced transitions of each participant, first reflecting the random transitions experienced during the training and single trials, then adapting to the first experienced graph structure and later to the second graph structure in sequence trials. In order to relate the SR models to participants’ response times, we calculated how surprising each transition in the sequence task was – assuming participants’ expectations were based on the current SR on the given trial, **M***^t^*. To this end, we normalized **M***^t^* to sum to 1, and then calculated the Shannon information (Shannon, 1948) for each trial, reflecting how surprising the just observed transition from stimulus *i* to *j* was given the history of previous transitions up to time point *t*:

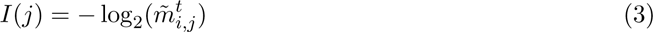

where 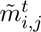 is the normalized (*i, j*)^th^ entry of SR matrix **M***^t^*. Rare events are more more surprising and require more information to represent them than common events. Using the base-2 logarithm allowed to express the units of information in bits (binary digits) and the negative sign ensured that the information measure was always positive or zero. In this case, Shannon information will be zero when the probability of an event is 1.0 or a certainty, i.e., there is no surprise (Shannon, 1948).

The final step in this analysis was to estimate LME models that tested how strongly this trial-wise measure of SR-based surprise was related to participants’ response times in the sequence task, for each level of the discount parameter *γ*. LME models therefore included fixed effects of the SR-based Shannon surprise, in addition to factors of task run, graph order (uni – bi vs. bi – uni) and graph structure (uni vs. bi) of the current run, as well as by-participant random intercepts and slopes. Separate LME models were conducted for each level of *γ* (twenty values for *γ* from 0 to 0.95 in steps of 0.05), and model comparison of the twenty models was performed using AIC, as reported in the main text. To independently investigate the effects of graph condition (uni vs. bi) and graph order (uni – bi vs. bi – uni), we analyzed separate LME models for each combination of the two factors, using only SR-based Shannon surprise as the main fixed effect of interest, and including by-participant random intercepts and slopes (see Fig. S5).

### Participant-specific SR model fitting

The above approach compared twenty LME models that were run across the data from all participants. In order to get an estimate of multi-step learning per participant, we performed model fitting to each participant’s data. We therefore fit the SR model to each participant’s data individually. Parameter fitting consisted of fitting only two parameters, namely the *α* and *γ* parameters of the SR model (see model description above). Model fitting minimized the negative log-likelihood of a GLM predicting response times (using an inverse gamma link function) within each participant using a nested approach akin to a coordinate descent approach (cf. Hall-McMaster et al., 2021; Koch et al., 2022). Specifically, the parameters *α* and *γ* were set in an outer loop using non-linear search method (NLOPT GN DIRECT L; Gablonsky and Kelley, 2001) implemented in the nloptr package (Johnson, 2019) in R. The GLM included a random intercept and regressors of the SR-based Shannon surprise, together with regressors for trial, task block, and the fingers used for responses. Note, that this modeling approach differed from the LME-based procedure described above, in that it used GLM with an inverse gamma link function and did not include additional regressors for trial and fingers used for the responses. The *β* coefficients of these regressors were then set using maximum likelihood estimation in an inner loop, and the resulting likelihood of the GLM was used to inform the non-linear search for the outer parameters (*α* and *γ*). Parameters were constrained to lie in the intervals given above. As shown in Fig. 3d, the best-fitting LME models of participants widely exhibited a significant relationship between Shannon surprise and response time on a trial, suggesting reliable estimates of the *γ* parameter. The value of the *γ* parameter directly influenced the Shannon surprise on each trial (see Eqns. 2 and 3). A non-significant relationship between response time and surprise (values above the dashed line) would indicate no effect on the LME model’s likelihood and suggest unreliable estimates of the *γ* parameter which controls the surprise on each trial.

After fitting the participant-specific parameters (*α* and *γ*) of the SR model for each participant, we used these parameters to generate the SR matrix for each participant at every trial during the task. To this end, we used the individually fitted parameters for *α* and *γ* as an input to Equation 2 to generate trial-by-trial SR matrices for every participant during the sequence task. Note, that the model parameters were fit based on a model that included all behavioral task data including data from the single trials (for details, see above), resulting in one value per parameter per participant. An example of the resulting SR matrices over the time course of the sequence task is shown in Fig. 3g.

### Analysis of classification accuracy and single trial classifier probability time courses

In order to assess the classifiers’ ability to differentiate between the neural activation patterns of individual visual objects and motor responses, we compared the predicted visual object or motor response of each example in the test set to the visual object or motor response that actually occurred on the corresponding trial. We obtained an average classification accuracy score for each participant by calculating the mean proportion of correct classifier predictions across all correctly answered single trials in session 1 (Fig. 4a). The mean decoding accuracy scores of all participants were then compared to the chance baseline of 100%*/*6 = 16.67% using a one-sided one-sample *t*-test, testing the a-priori hypothesis that mean classification accuracy would be higher than the chance baseline. The effect size (Cohen’s *d*) was calculated as the difference between the mean of accuracy scores and the chance baseline, divided by the standard deviation of the data (Cohen, 1988). These calculations were performed separately for each ROI and the resulting *p*-values were adjusted for multiple comparisons using Bonferroni correction (Bonferroni, 1936).

We also examined the effect of task run on classification accuracy in single trials. To this end, we conducted an LME model including the task run as the main fixed effect of interest as well as by-participant random intercepts and slopes. We then assessed whether performance was above the chance level for all nine task runs and conducted nine separate one-sided one-sample *t*-tests separately per ROIs, testing the a-priori hypothesis that mean decoding accuracy would be higher than the 16.67% chance-level in each task run. All *p*-values were adjusted for 18 multiple comparisons (across nine runs and two ROIs) using the Bonferroni-correction (Bonferroni, 1936).

Furthermore, we assessed the classifiers’ ability to accurately detect the presence of visual objects and motor responses on single trials. For this analysis we applied the trained classifiers to fifteen volumes from the volume closest to the event onset and examined the time courses of the probabilistic classification evidence in response to the event on single trials (Fig. 4b). In order to test if the time series of classifier probabilities reflected the expected increase of classifier probability for the event occurring on a given trial, we compared the time series of classifier probabilities related to the classified class with the mean time courses of all other classes using a two-sided paired *t*-test at the fourth TR from event onset. Classifier probabilities were normalized by dividing each classifier probability by the sum of the classifier probabilities across all fifteen TRs of a given trial. Here, we used the Bonferroni-correction method (Bonferroni, 1936) to adjust for multiple comparisons of two observations (one test per ROI). In the main text, we report the results for the peak in classification probability of the true class, corresponding to the fourth TR after stimulus onset. The effect size (Cohen’s *d*) was calculated as the difference between the means of the probabilities of the current versus all other stimuli, divided by the standard deviation of the difference (Cohen, 1988).

### Modeling of stimulus-driven classifier time courses in on-task intervals

In our previous work (Wittkuhn and Schuck, 2021), we showed that analyzing probabilistic classifier evidence on single trials of a presented stimulus revealed multivariate decoding time courses that can be characterized by a slow response function that resembles single-voxel hemodynamics. Here, we applied the same methodology to capture the expected effects of stimulus-driven activity elicited by previous trials in on-task intervals. The details of this modeling approach were first described in Wittkuhn and Schuck (2021), but, for completeness, we reproduce them here as well. Specifically, we modeled an individual classifier probability response function as a sine wave that was flattened after one cycle, scaled by an amplitude and adjusted to baseline. The model was specified as follows:

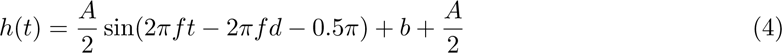

where *A* is the response amplitude (the peak deviation of the function from baseline), *f* is the angular frequency (unit: 1/TR, i.e., 1*/*1.25 = 0.8 Hz), *d* is the onset delay (in TRs), and *b* is the baseline (in %). The restriction to one cycle was achieved by converting the sine wave in accordance with the following piecewise function:

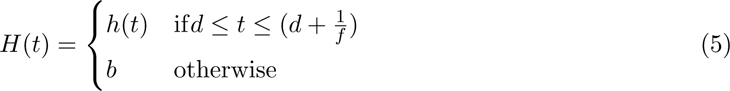

As in Wittkuhn and Schuck (2021), we fitted the four model parameters (*A*, *f*, *d* and *b*) to the mean probabilistic classifier evidence of each stimulus class at every TR separately for each participant. For convenience, we count time *t* in TRs. The default parameters (as well as lower and upper bounds) were set to *A* = 0.6 (0.1 *≤ A ≤* 1.0), *f* = 0.2 (0.01 *≤ f ≤* 0.5), *d* = 0.0 (0 *≤ d ≤* 8), and *b* = 0.1 (0.0 *≤ b ≤* 0.3), respectively. The parameters were optimized using a version of the Constrained Optimization BY Linear Approximations (COBYLA) algorithm for derivative-free optimization with nonlinear inequality and equality constraints (NLOPT LN COBYLA; Powell, 1994) implemented in the nloptr package (Johnson, 2019) in R. The relative tolerance for convergence was set to 1.0 *×* 10*^−^*^8^. The maximum number of evaluations allowed during optimization was set to 1.0 *×* 10^5^. This function was then fit to the mean classifier time courses across fifteen TRs on single trials, separately for each stimulus class and participant.

To assess the accuracy of the sine-based response function in predicting observed data, we implemented an evaluation function, aiming to fine-tune model parameters for improved predictive performance. Specifically, the sum of squared errors (SSE) was computed to quantify the disparity between the predicted values (y) and the observed data (see Equation 6). The SSE metric serves as the indicator of the model’s fidelity to the observed data, with lower values indicative of a more accurate fit. The SSE output from this evaluation function guides our optimization algorithm in iteratively adjusting model parameters to minimize the disparity between predicted and observed data, ultimately refining the accuracy of the sine-based response model.

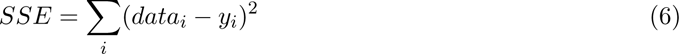

Next, we modeled stimulus-evoked classifier probabilities lagging into the on-task interval. To this end, we first took the four model parameters (*A*, *f*, *d* and *b*) that were fit individually to the classifier probability data on single trials of each participant, ROI and stimulus and convolved them with the onsets of ten stimuli *before* and ten stimuli *after* the on-task intervals in the sequence task. Given the timings of the trial procedure (see Methods above), we reasoned that considering ten stimuli before the on-task interval would sufficiently capture any stimulus-evoked activity leaking into the on-task interval. Specifically, with a fixed stimulus duration of 800 ms and an average ITI of 750 ms (for details, see the description of the trial procedure in the Methods above) this would amount to considering stimulus-driven activity of up to 15.5 s on average before the on-task interval, which roughly amounts to the expected duration of the canonical HRF (see e.g., Aguirre et al., 1998). We reasoned that additionally considering stimuli *after* the on-task interval would allow to account for and investigate any combination of stimulus-evoked and replay-driven activity that might extend beyond the on-task intervals. Of note, due to trial-specific timing of task components, not all trials had all stimuli occur before them, i.e., for some trials some stimuli never occurred in the ten preceding trials.

Next, we calculated the mean stimulus-evoked activity for each TR of the on-task interval separately for each participant. Note, that the sine-based response model is a continuous function that can be evaluated over any arbitrarily densely sampled interval. Of note, the average value of a continuous function on an interval *f*_avg_ can be approximated by a list of points to derive that the average value is proportional to the area under the curve, i.e., the definite integral:

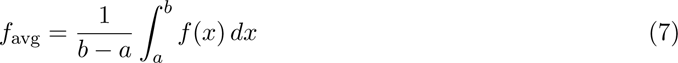

Here, as a result of our model fitting procedure (see above) we evaluated the function across a time window of 10 TRs with a constant sampling frequency of 0.1. Note, that increasing the sampling frequency (e.g., to 0.01 or 0.001) would only marginally improve the approximation of the mean of the continuous function within a certain interval at the expense of increased computation time. As the same stimulus could occur multiple times on the interval-preceding trials, this might result in overlapping activation of the same stimulus. To account for overlapping activation from the same stimulus, we summed the modeled activation values separately for each stimulus at each TR. To obtain one activation value per TR, we considered all data points that would fall within the range of one TR of 1.25 s. When a specific TR did not contain any stimulus-related activity according to the modeling approach, the activation of that stimulus at that time point was set to 0. An illustration of the resulting modeled stimulus-driven classifier probability time courses is shown in Fig. S10.

### Modeling of classifier time courses in on-task intervals

After generating predictions for stimulus-driven classifier probabilities during on-task intervals, we conducted analyses of classifier probabilities related to putative replay events using LME models while accounting for stimulus-evoked activity. Specifically, we compared four different models in a model comparison. All models accounted for individual variability by including a by-participant random intercept term. Random by-participant slopes were removed in order to achieve a non-singular fit of the model (for details on statistical analyses, see above). First, a baseline model (Model 1) described the relationship between classifier probabilities (as the main response variable) and modeled stimulus-evoked activity as the fixed effect predictor. All subsequent model specifications extended this baseline model by one or more additional fixed effect predictor variables. Specifically, we conceived an additional LME that investigated the influence of the one-step task transition probabilities (onestep model; Model 2). In addition, we investigated the influence of the probability of individual task nodes in the SR model to the respective time points of the on-task intervals, either as the sole predictor variable (multi-step SR model; Model 3) or in combination with the predictor of one-step task transition probabilities (one-step + multi-step SR model; Model 4). To obtain the SR probability of each node at each time point of the on-task interval, we extracted these values from the continuously updated SR matrices of each participants (for details, see above). The models were fit to all data of the sequence learning task, separately for each ROI and TR. This procedure then allowed to compare the models at each TR of on-task intervals, quantified by their AIC scores.

### Statistical analyses of classifier time courses on sequence trials

Classifier probabilities on sequence trials indicated that the fMRI signal was strongly dominated by the activation of the event on the current trial. In order to test this effect, we calculated the mean classifier probabilities for the current and all other five events of the current trial across all eight TRs in the ITIs. The mean classifier probabilities of the current event were then compared to the mean classifier probabilities of all other events using two two-sided paired *t*-tests, one for each ROI. The Bonferroni-correction method Bonferroni (1936) was used to correct the *p*-values for two comparisons. The effect size (Cohen’s *d*) was calculated as the difference between the means of the probabilities of the current versus all other events, divided by the standard deviation of the difference Cohen (1988).

### Calculating the TR-wise sequentiality metric

To analyze evidence for sequential replay during on-task intervals in sequence trials, we calculated a sequentiality metric quantified by the slope of a linear regression between the classifier probabilities or model residuals and the sequential orderings of a 5-item sequence in each TR, similar to our previous work (Wittkuhn and Schuck, 2021). The mean slope coefficients of all participants were then compared to zero at each TR using a series of two-sided one-sample *t*-test, separately for each condition (depending on the analyses, separated by ROI, graph structure and / or conscious knowledge). *p*-values were adjusted for multiple comparisons using FDR or Bonferroni correction (Bonferroni, 1936), as reported in the main text. The effect size (Cohen’s *d*) was calculated as the difference between the mean of slope coefficients and the baseline, divided by the standard deviation of the data (Cohen, 1988).

## Data and code availability statement

All code and data used in this study will be made available upon publication in a peer-reviewed journal.

## Acknowledgments

This work was supported by an Independent Max Planck Research Group grant awarded to N.W.S by the Max Planck Society (M.TN.A.BILD0004), a Starting Grant awarded to N.W.S by the European Union (ERC-2019-StG REPLAY-852669) and funding by the Federal Ministry of Education and Research (BMBF) and the Free and Hanseatic City of Hamburg under the Excellence Strategy of the Federal Government and the Länder awarded to N.W.S. L.W. was a pre-doctoral fellow of the International Max Planck Research School on Computational Methods in Psychiatry and Ageing Research (IMPRS COMP2PSYCH). The participating institutions are the Max Planck Institute for Human Development, Berlin, Germany, and University College London, London, UK. For more information, see https://www.mps-ucl-centre.mpg.de/en/comp2psych. We also thank Leonardo Pettini for help with task development, Gregor Caregnato for help with participant recruitment and study coordination, Sonali Beckmann, Sam Chien (https://orcid.org/0000-0003-4306-1308), Theresa Fox https://orcid.org/0000-0002-7345-6354, Sam Hall-McMaster (https://orcid.org/0000-0003-1641-979X), Nir Moneta (https://orcid.org/0000-0001-6125-4117), Liliana Polanski (https://orcid.org/0000-0002-0842-8787), Nadine Taube, and Kateryna Yasynska – in alphabetical order of last names – for assistance with MRI data acquisition, Anika Löwe (https://orcid.org/0000-0003-3132-5767) for help with MRI data collection and comments on a previous version of this manuscript, Onďrej Źıka (https://orcid.org/0000-0003-0483-4443) for help with MRI data collection and statistical analyses, Michael Krause for help with high performance computing (HPC), all members of the Max Planck Research Group NeuroCode for helpful discussions about the contents of this manuscript, and all participants for their participation. For a visual depiction of the true team effort during data collection, see Fig. S13.

## Author Contributions

The following list of author contributions is based on the CRediT taxonomy (Brand et al., 2015). For details on each type of author contribution, see Brand et al. (2015).

Conceptualization: L.W., N.W.S.; Methodology: L.W., C.K., L.M.K., N.W.S.; Software: L.W., C.K., L.M.K., N.W.S.; Validation: L.W.; Formal analysis: L.W., C.K., N.W.S.; Investigation: L.W., L.M.K.; Resources: L.W., N.W.S.; Data curation: L.W., L.M.K.; Writing - original draft: L.W.; Writing - review & editing: L.W., C.K., L.M.K., N.W.S.; Visualization: L.W., C.K.; Supervision: N.W.S.; Project administration: L.W., N.W.S.; Funding acquisition: N.W.S.

## Glossary

AC-PC: anterior commissure - posterior commissure.
AIC: Akaike information criterion.
ANOVA: analysis of variance.
ANTs: Advanced Normalization Tools.
A-P: anterior-to-posterior.
BIDS: brain imaging data structure.
BOBYQA: Bound Optimization BY Quadratic Approximation.
BOLD: blood-oxygen-level dependent.
CI: confidence interval.
COBYLA: Constrained Optimization BY Linear Approximations.
CSF: cerebrospinal fluid.
DGPs: German Psychological Society.
DICOM: Digital Imaging and Communications in Medicine.
EPI: echo-planar imaging.
FA: flip angle.
FD: framewise displacement.
FDR: false discovery rate.
fMRI: functional magnetic resonance imaging.
FOV: field of view.
FSL: FMRIB Software Library.
FWHM: Full Width at Half Maximum.
GLM: general linear model.
GM: gray-matter.
GR: gradient recalled.
HPC: high performance computing.
HRF: The hemodynamic response function (HRF) characterizes an fMRI response that results from a brief, spatially localized pulse of neuronal activity.
HSD: honest significant difference.
INU: intensity non-uniformity.
IQR: interquartile range.
ITI: inter-trial interval.
LME: linear mixed effects.
LTS: long-term support.
MB: multi-band.
min: minute.
MPRAGE: Magnetization Prepared Rapid Gradient Echo.
MRI: magnetic resonance imaging.
ms: millisecond.
PFC: prefrontal cortex.
PMU: Physiological Measurement Unit.
ROI: region of interest.
s: second.
SD: standard deviation.
SEM: standard error of the mean.
SI: supplementary information.
SR: successor representation.
S-R: stimulus-response.
SRI: stimulus-response interval.
SSE: sum of squared errors.
SUSAN: Smallest Univalue Segment Assimilating Nucleus.
T1w: T1-weighted.
TD: temporal difference.
TE: echo time.
TI: inversion time.
TR: repetition time.
WM: white-matter.

## Supplementary Information

### Supplementary Figures

**Supplementary Figure S1:**
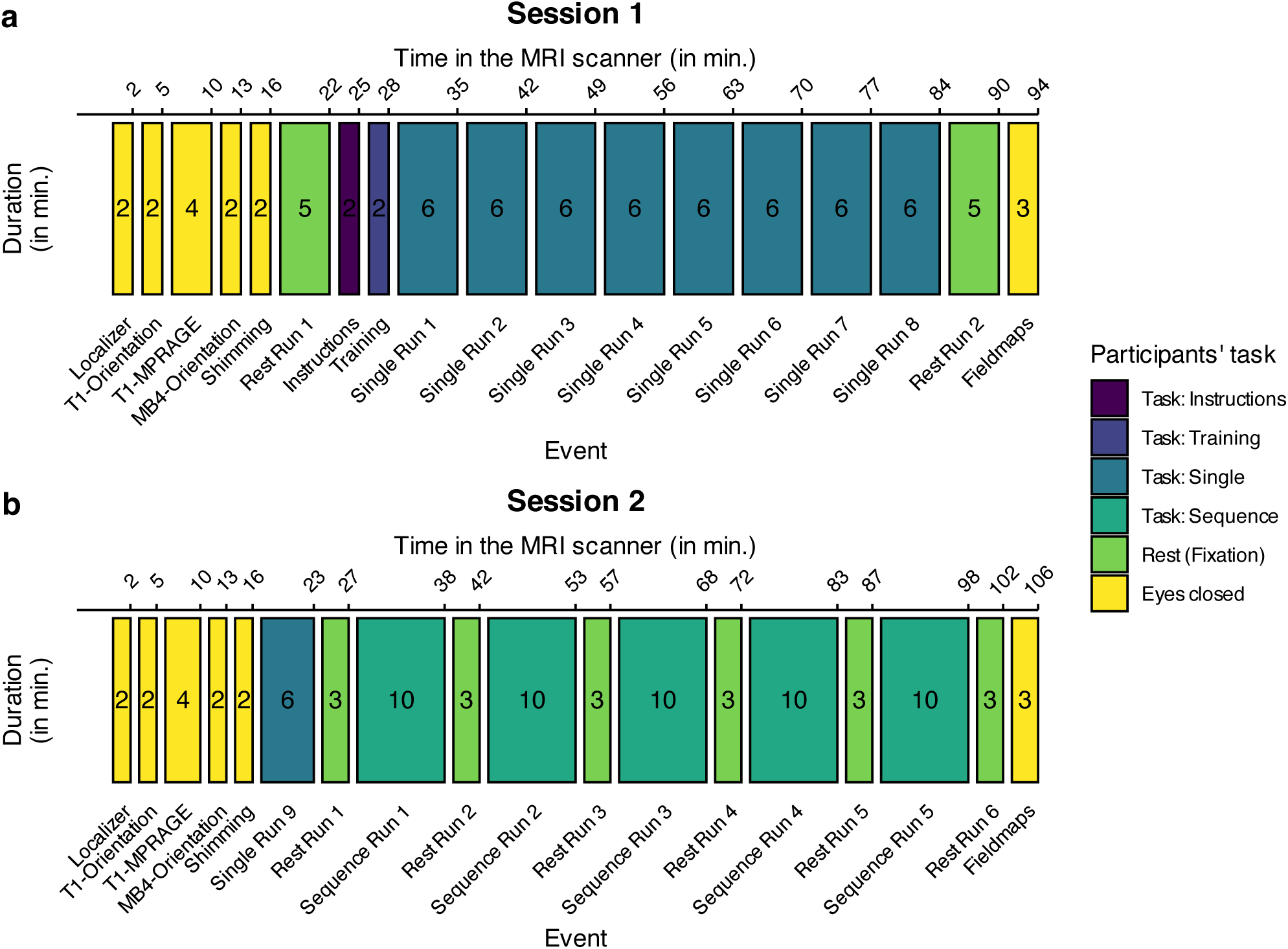
Study procedure. **(a)** Session 1 started with a 5 minutes (mins) resting-state scan before participants read the task instructions and completed the training condition of the task. Participants then completed eight runs of the single trial condition of about 6 min each before another 5 min resting-state scan was recorded. **(b)** Session 2 started with another run of the single trial condition of about 6 min. Participants then completed all five runs of the sequence task of about 10 min each which were interleaved with six resting-state scans of 3 min each. Both experimental sessions started with a short localizer scan, a T1w anatomical MRI scan as well as advanced shimming and ended with the recording of fieldmaps (for details on MRI acquisition, see Methods). Participants were asked to keep their eyes closed during these scans and other additional preparations by the study staff, e.g., orientation of the field of view (FOV). The numbers inside the rectangles indicate approximate duration of each procedure in min. Colors indicate participants’ task during each study event (see legend). All timings are rough estimates based on the task design and may have varied within and between individual participants.

**Supplementary Figure S2:**
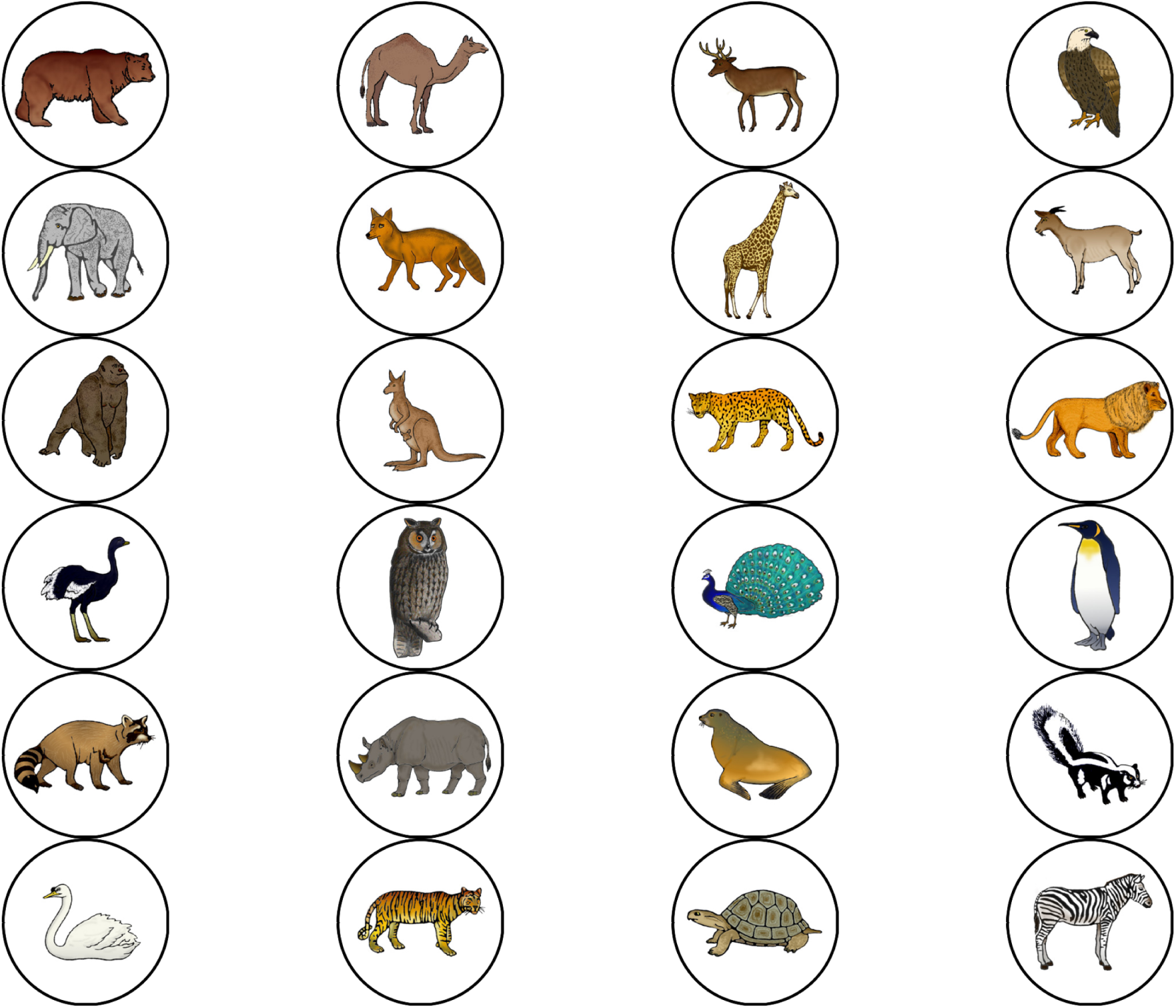
Overview of all. 24 **animal stimuli.** All visual stimuli were taken from a set of colored and shaded images commissioned by Rossion and Pourtois (2004), which are loosely based on images from the original Snodgrass and Vanderwart set (Snodgrass and Vanderwart, 1980). The images are freely available from the internet at https://sites.google.com/andrew.cmu.edu/tarrlab/stimuli under the terms of the Creative Commons Attribution-NonCommercial-ShareAlike 3.0 Unported license (CC BY-NC-SA 3.0; for details, see https://creativecommons.org/ licenses/by-nc-sa/3.0/) and have been used in similar previous studies (e.g., Garvert et al., 2017). Stimulus images courtesy of Michael J. Tarr, Carnegie Mellon University, (for details, see http://www.tarrlab.org/). In total, we selected 24 images which depicted animals that could be expected in a public zoo. Specifically, the images depicted a bear, a dromedary, a deer, an eagle, an elephant, a fox, a giraffe, a goat, a gorilla, a kangaroo, a leopard, a lion, an ostrich, an owl, a peacock, a penguin, a raccoon, a rhinoceros, a seal, a skunk, a swan, a tiger, a turtle, and a zebra (in alphabetical order, from left to right and top to bottom). For each participant, six task stimuli were randomly selected from the set of 24 the animal images and each image was randomly assigned to one of six response buttons. For more details, see the task description in the Methods section.

**Supplementary Figure S3:**
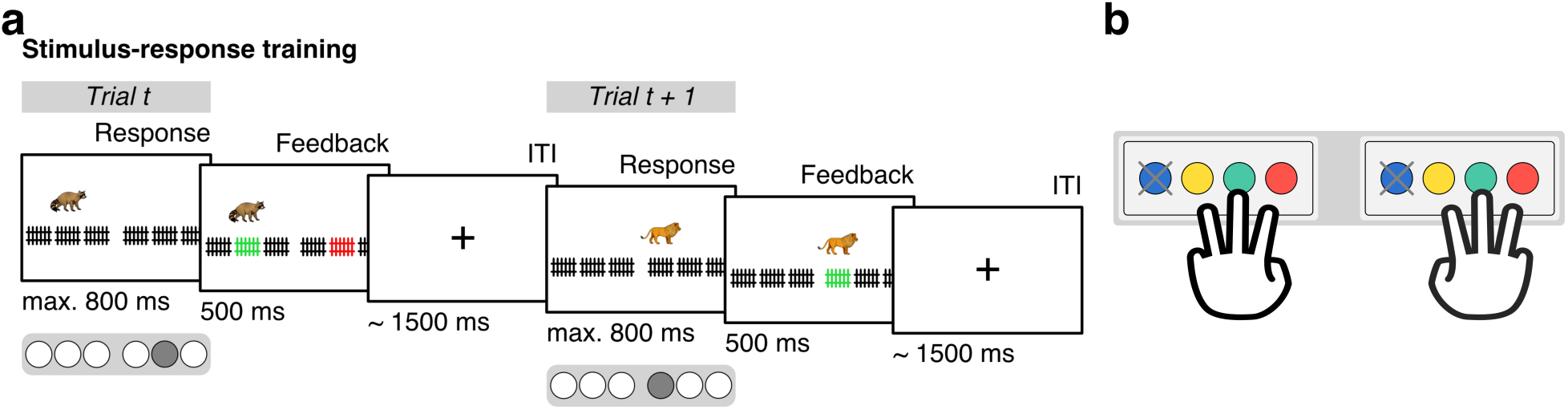
Stimulus-response (S-R) training and response mapping. **(a)** On training trials, participants were explicitly trained to learn associations between six animal stimuli and six response buttons. On each trial, an animal appeared above one of six cages that were representing the six response buttons. Participants were asked to press the corresponding response button within 800 ms and subsequently received feedback if their response was incorrect (see *Trial t*) or correct (see *Trial t + 1*). fMRI data from training trials were not used for any further analysis. For more details, see the task description in the Methods section. **(b)** Hand placement during behavioral responses in the MRI scanner. Each of the six animals was associated with one of six response buttons that participants controlled with their index, middle, and ring finger of both hands. The response pad included a fourth button that participants were asked to ignore. Colors indicate the actual color of each button and are not related to any color in the task or figures in the main text. The illustration of the hands in (b) is an artwork “three fingers” by Herbert Spencer from the Noun Project, licensed under Creative Commons Attribution 3.0 United States (CC BY 3.0 US; for details, see https://creativecommons.org/licenses/by/3.0/us/) and available from https://thenounproject.com/term/three-fingers/155721/. All visual stimuli in (a) were taken from a set of colored and shaded images commissioned by Rossion and Pourtois (2004), which are loosely based on images from the original Snodgrass and Vanderwart set (Snodgrass and Vanderwart, 1980). The images are freely available from the internet at https://sites.google.com/andrew.cmu. edu/tarrlab/stimuli under the terms of the Creative Commons Attribution-NonCommercial-ShareAlike 3.0 Unported license (CC BY-NC-SA 3.0; for details, see https://creativecommons.org/licenses/by-nc-sa/3.0/) and have been used in similar previous studies (e.g., Garvert et al., 2017). Stimulus images courtesy of Michael J. Tarr, Carnegie Mellon University, (for details, see http://www.tarrlab.org/).

**Supplementary Figure S4:**
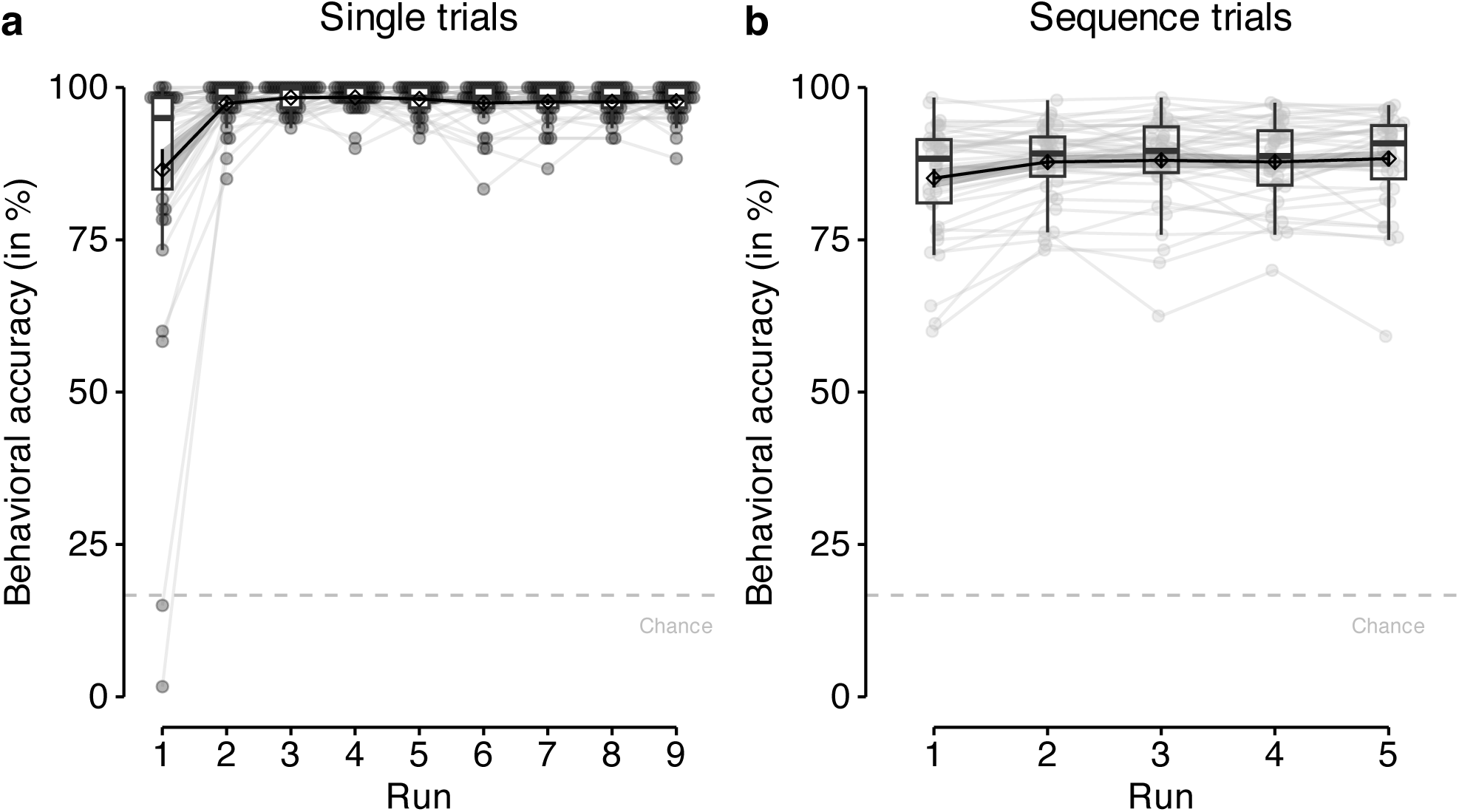
Behavioral accuracy per task run in single and sequence trials. **(a)** Mean behavioral accuracy (in %; y-axis) per run (x-axis) in single trials. **(b)** Mean behavioral accuracy (in %; y-axis) per run (x-axis) in sequence trials. The chance level (gray dashed line) is at 16.67%. Each gray dot corresponds to averaged data from one participant. Gray lines connect data across runs for each participant. Boxplots indicate the median and IQR. The lower and upper hinges correspond to the first and third quartiles (the 25^th^ and 75^th^ percentiles). The upper whisker extends from the hinge to the largest value no further than 1.5*∗* IQR from the hinge (where IQR is the interquartile range (IQR), or distance between the first and third quartiles). The lower whisker extends from the hinge to the smallest value at most 1.5*∗* IQR of the hinge. The diamond shapes show the sample mean. Error bars and shaded areas indicate *±*1 SEM. All statistics have been derived from data of *n* = 39 human participants who participated in one experiment.

**Supplementary Figure S5:**
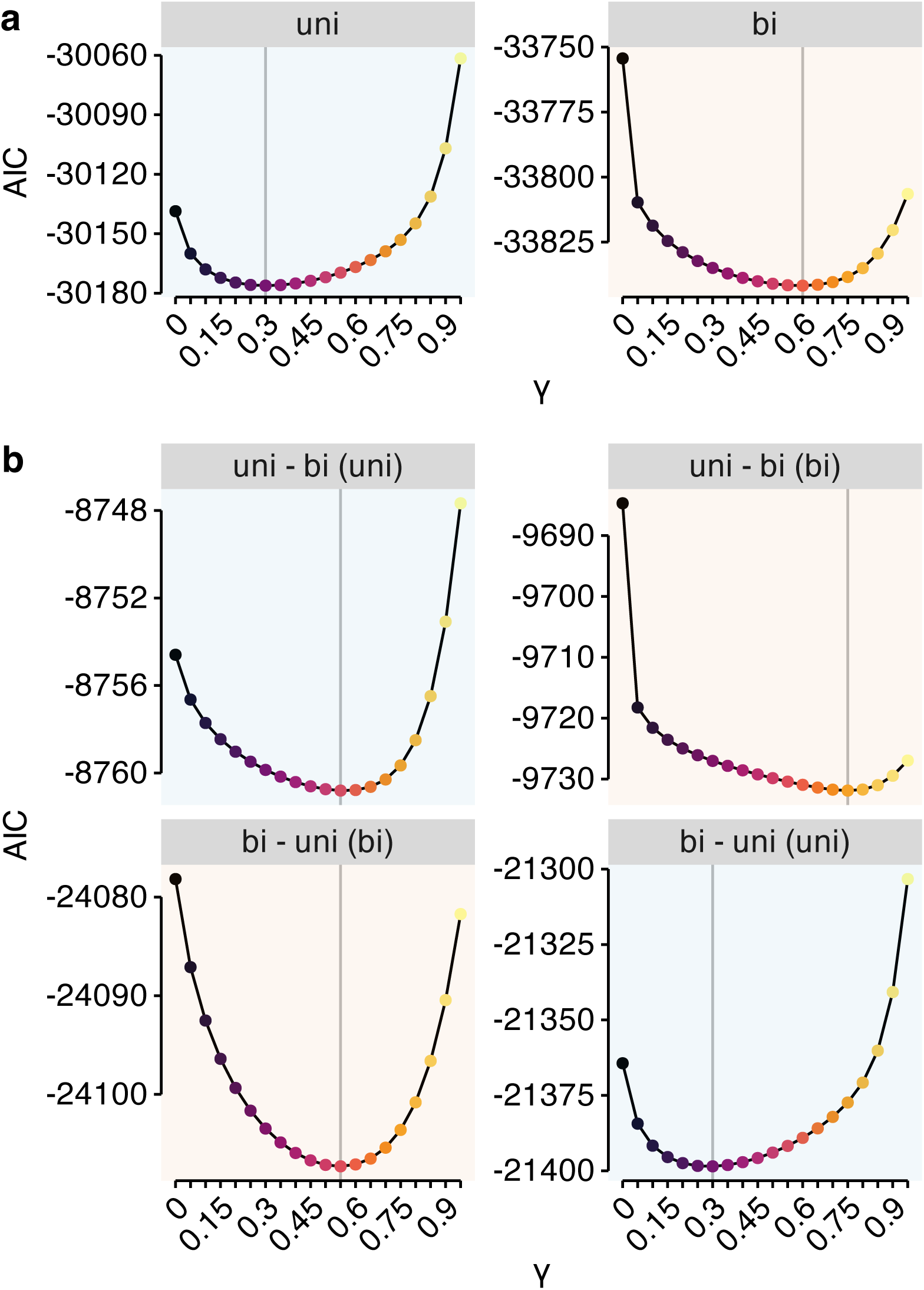
Influence of graph condition and graph order on SR-based modeling of response time. **(a)** AIC scores (y-axis) for LME models fit to participants’ log response time data using Shannon information based on SRs with varying predictive horizons (the discounting parameter *γ*; x-axis) as the predictor variable, separated by graph condition (uni vs. bi). **(b)** AIC scores (y-axis) for LME models fit to participants’ log response time data using Shannon information based on SRs with varying predictive horizons (the discounting parameter *γ*; x-axis) as the predictor variable, separated by graph order (uni – bi vs. bi – uni; horizontal panels) and graph condition (uni vs. bi; panel colors). Vertical lines in (a) and (b) mark the lowest AIC score. All statistics have been derived from data of *n* = 39 human participants who participated in one experiment.

**Supplementary Figure S6:**
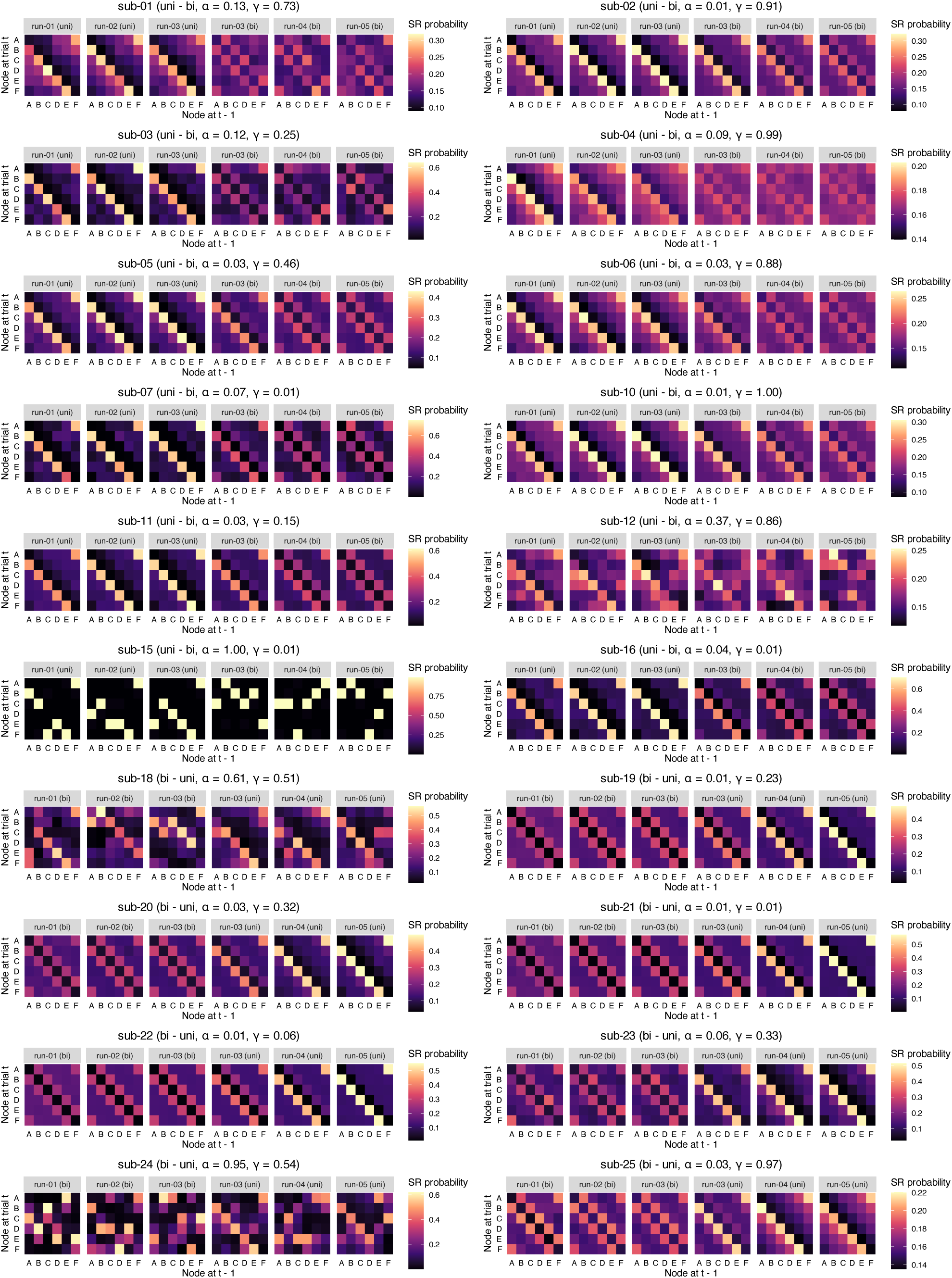
[for a caption, see caption of Fig. 3g]

**Supplementary Figure S7:**
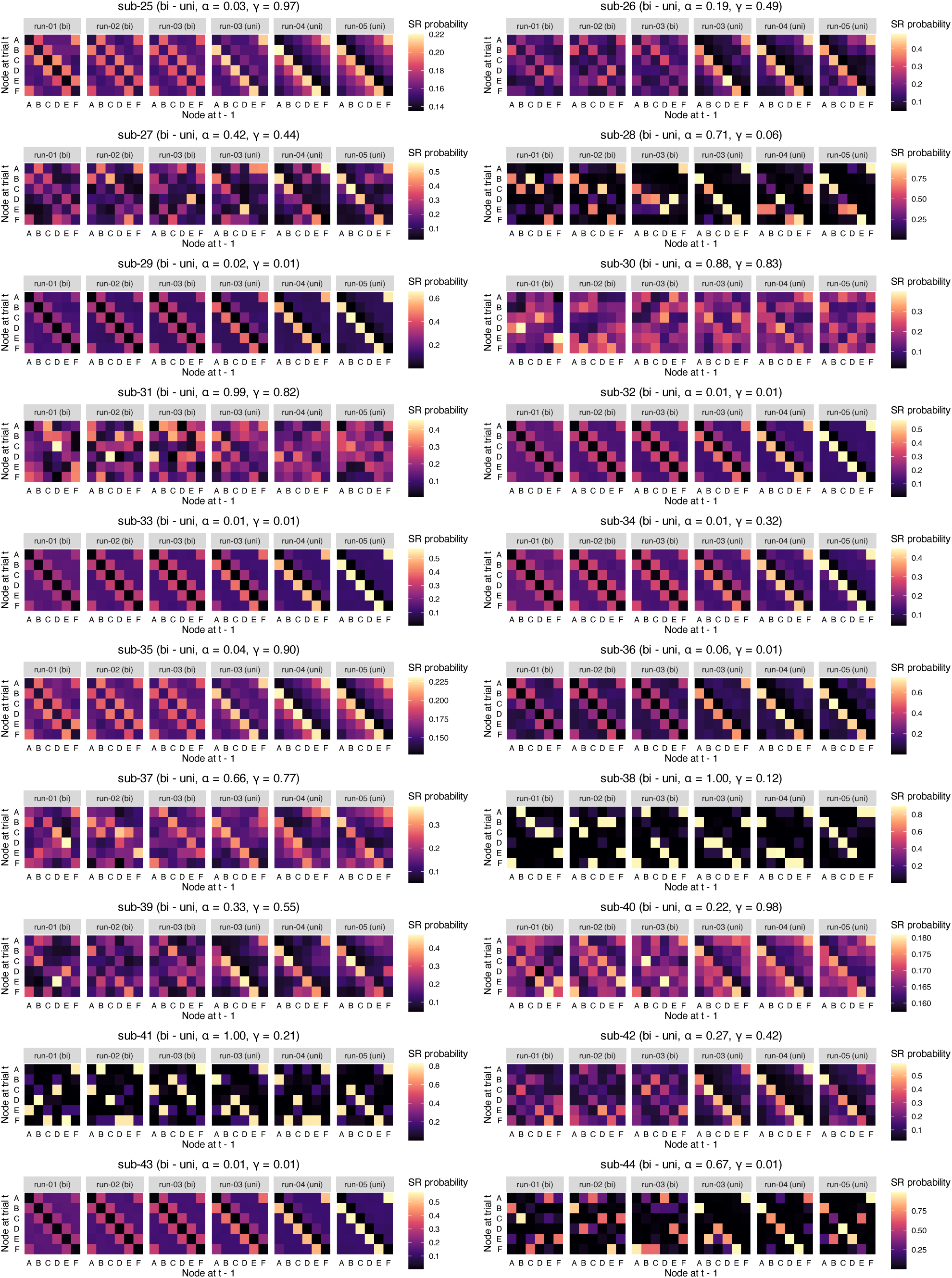
[for a caption, see caption of Fig. 3g]

**Supplementary Figure S8:**
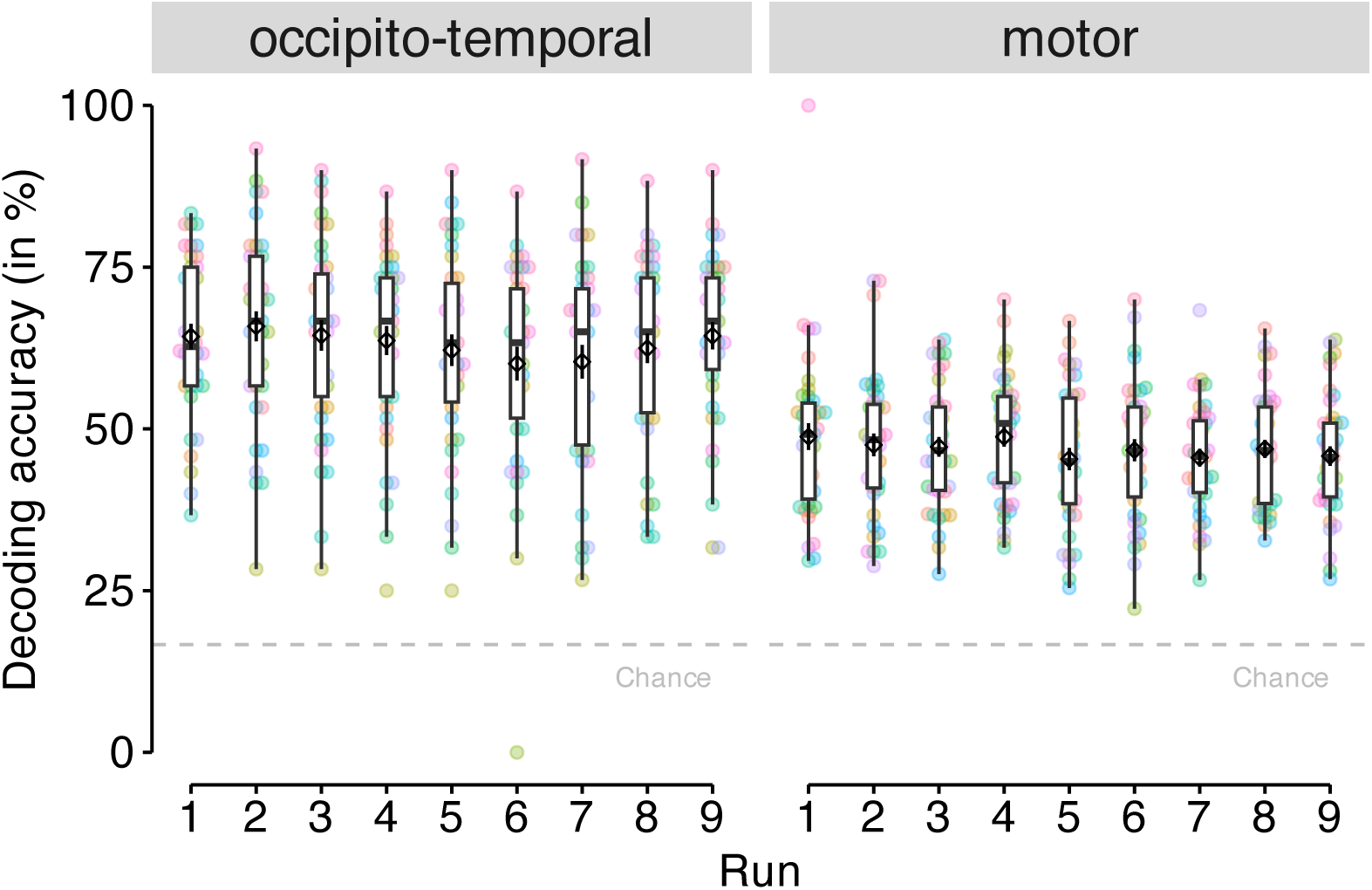
Classification accuracy for each run of single trials. Cross-validated classification accuracy (in %) in decoding six unique visual objects in occipito-temporal data (left panel) and six unique motor responses in sensorimotor cortex data (right panel) during task performance on single trials, separately for each run (x-axis). Chance level for each run is at 16.67% (horizontal dashed line). Boxplots indicate the median and IQR. The lower and upper hinges correspond to the first and third quartiles (the 25*^th^* and 75*^th^* percentiles). The upper whisker extends from the hinge to the largest value no further than 1.5*∗* IQR from the hinge (where IQR is the interquartile range (IQR), or distance between the first and third quartiles). The lower whisker extends from the hinge to the smallest value at most 1.5*∗* IQR of the hinge. The diamond shape show the sample mean. Error bars indicate *±*1 SEM. Each dot corresponds to averaged data from one participant. All statistics have been derived from data of *n* = 39 human participants who participated in one experiment.

**Supplementary Figure S9:**
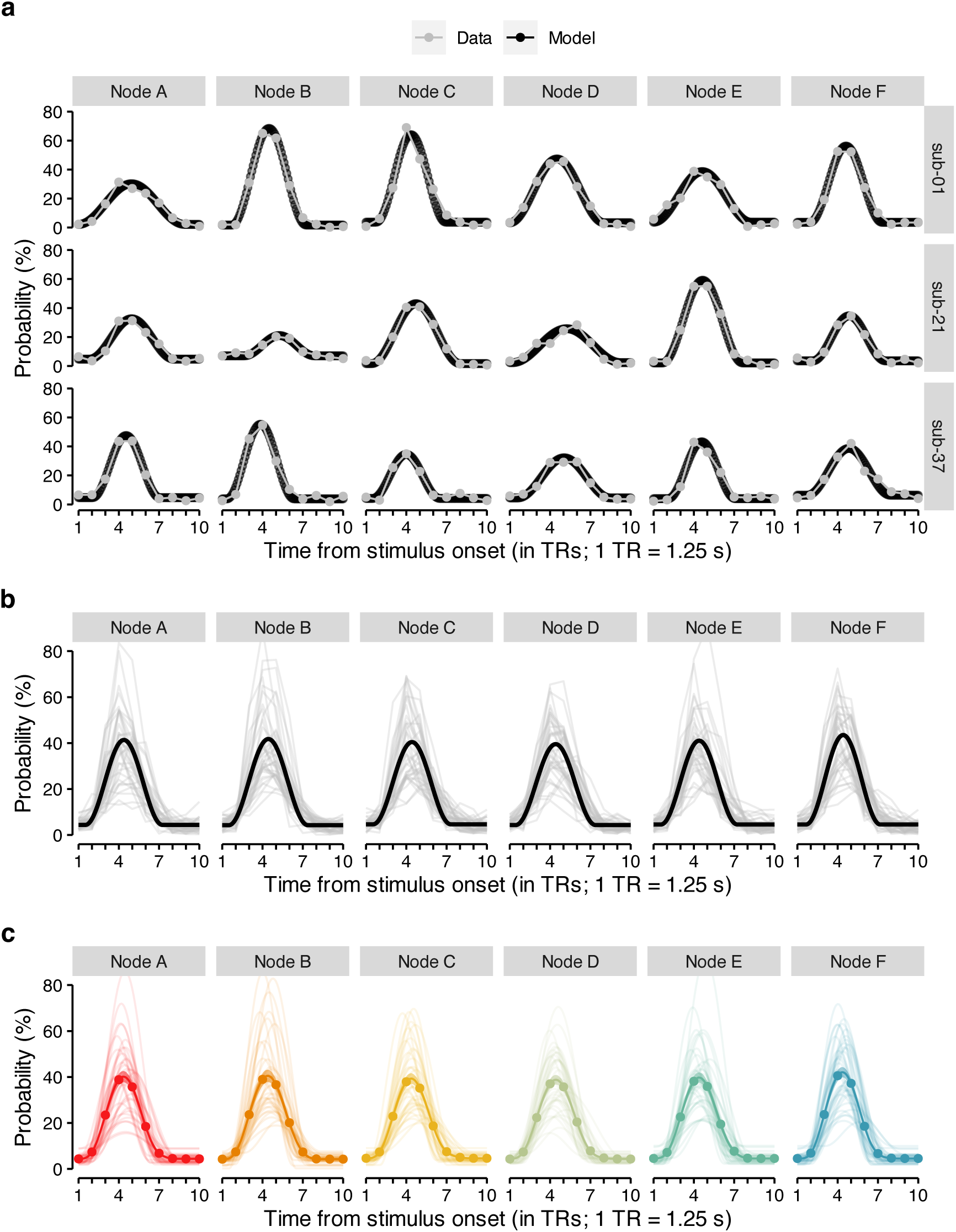
Individual fits of sine wave response function to probabilistic classifier evidence. **(a)** Time courses (in TRs from stimulus onset; x-axis) of probabilistic classifier evidence (in %; y-axis) generated by the sine wave response function with fitted parameters (black dotted line) or the true data (gray line and dots) separately for the six stimulus classes (vertical panels) and three randomly chosen example participants (horizontal panels). **(b)** Time courses (in TRs from stimulus onset; x-axis) of mean probabilistic classifier evidence (in %; y-axis) averaged separately for each participant (gray semi-transparent lines) and each of the six stimulus class (vertical panels) or predicted by the sine wave response model based on fitted parameters averaged across all participants (black line). **(c)** Time courses (in TRs from stimulus onset; x-axis) of mean probabilistic classifier evidence (in %; y-axis) as predicted by the sine wave response model based on fitted parameters derived separately for each participant (individual, semi-transparent lines). Each semi-transparent line in (b) and (c) represents data from one participant. Classifier probabilities in (a), (b) and (c) were normalized across 15 TRs. The chance level therefore is at 100*/*15 = 6.67%. 1 TR = 1.25 s. All statistics have been derived from data of *n* = 39 human participants who participated in one experiment.

**Supplementary Figure S10:**
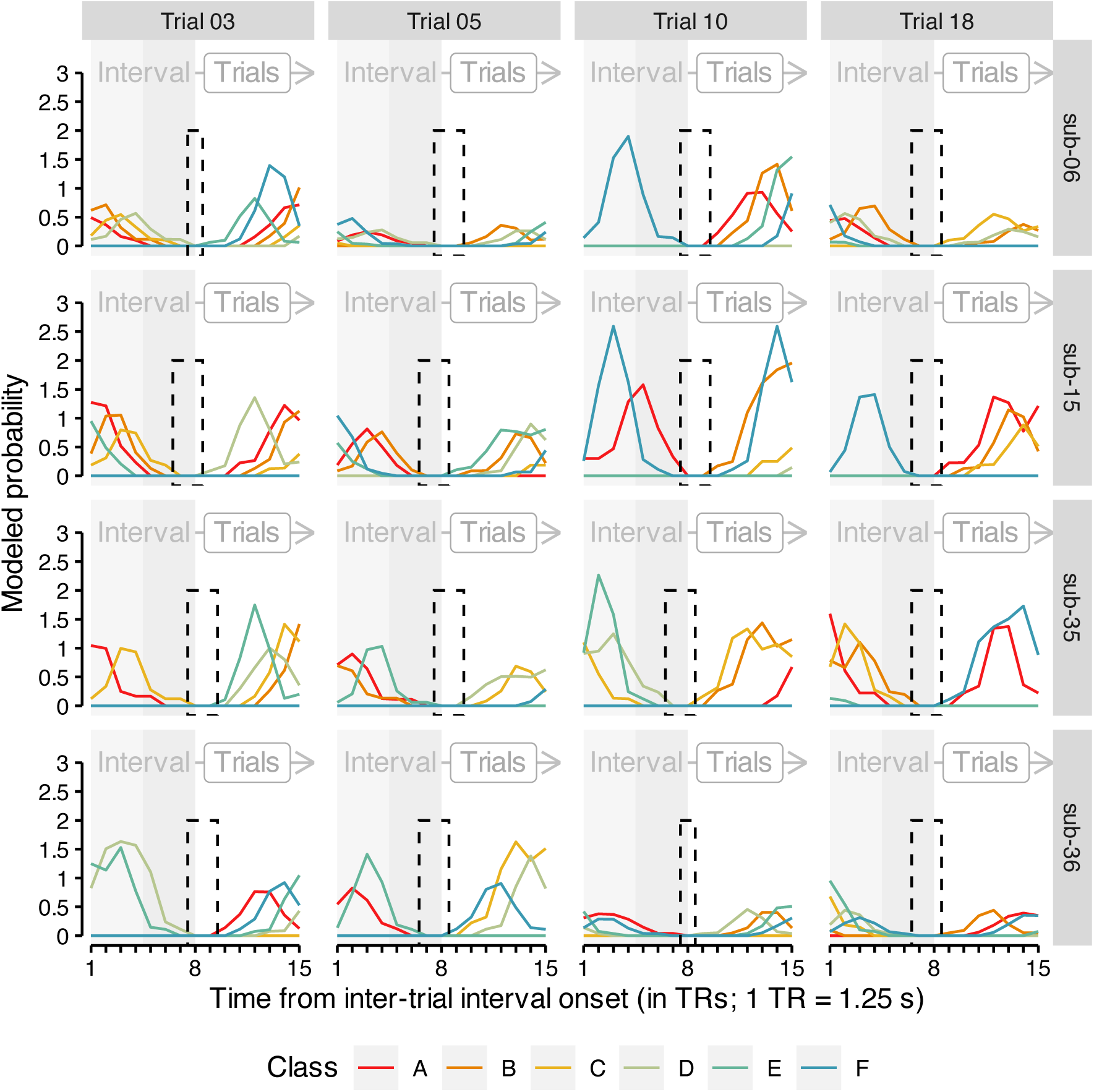
Illustration of modeled stimulus-driven activity during on-task intervals. Time courses (in TRs from inter-trial interval (ITI) onset; x-axis; light and dark gray background) of modeled probabilistic classifier evidence (in %; y-axis) based on the sine wave response function with individually fitted parameters separately for the six stimulus classes (colors; see legend), four randomly chosen example participants (horizontal panels) and four randomly chosen example trials (vertical panels). Each line in represents data for one stimulus class from one participant on a particular trial. Rectangles with dashed lines illustrate TR intervals with no expected stimulus-driven activity (based on the modeling approach). 1 TR = 1.25 s. Source data are provided as a Source Data file.

**Supplementary Figure S11:**
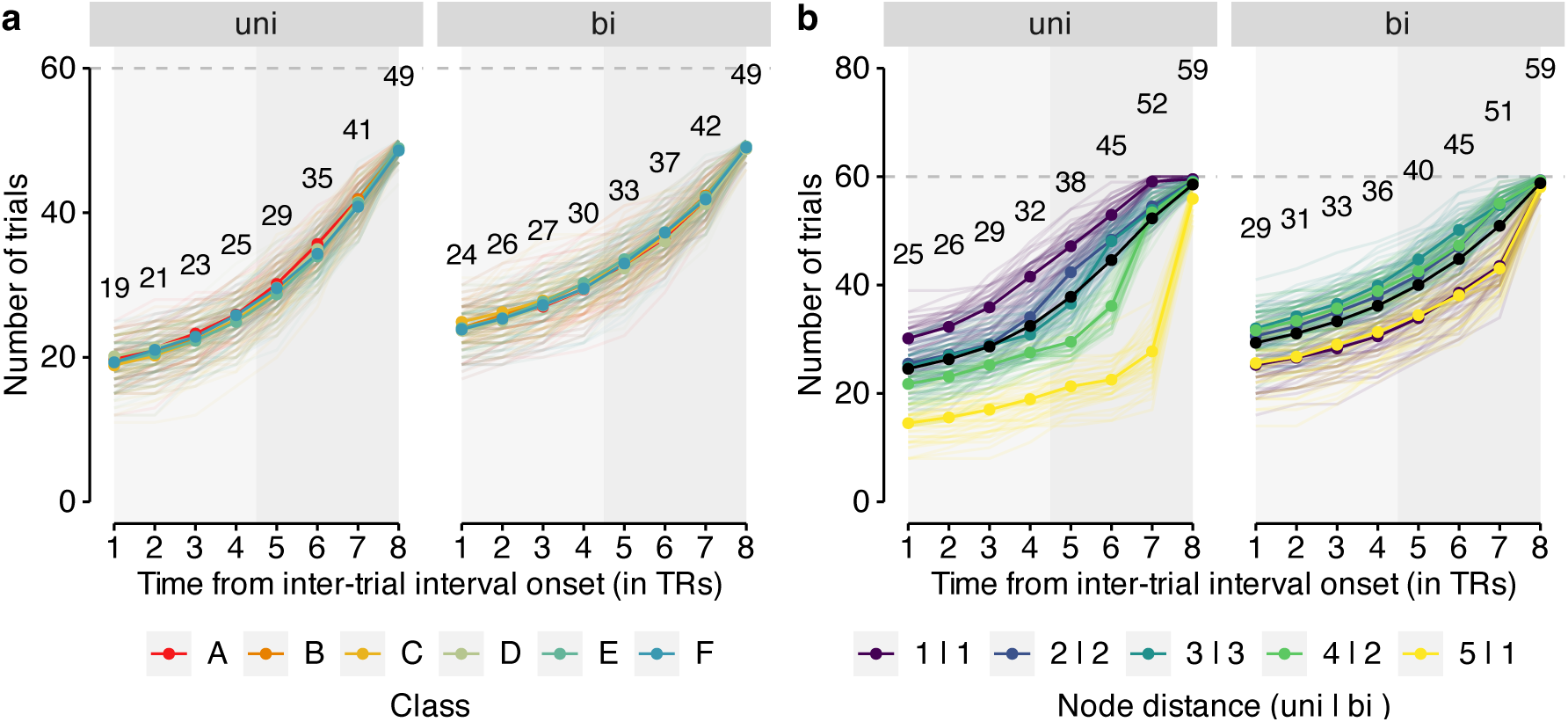
Analysis of classifier probabilities in subsets of sequence trials after removing data points expected to contain stimulus-evoked activity. **(a)** Number of trials (y-axis) for each TR from intertrial- interval (ITI) onset (x-axis; light and dark gray background) separately for each of the six stimulus classes (colors; see legend) and both graph structures (uni and bi; vertical panels). **(b)** As in (a), but for each graph node distance. The black line represents the mean number of trials across classes of node distances. The maximum number of trials for each stimulus class (a) or node distance (b) is 60 (horizontal dashed gray line). Each semi-transparent line represents data from one participant. Numbers indicate mean number of trials across all participants and stimulus classes (a) or node distances (b). All data are from the occipito-temporal anatomical ROIs. Qualitatively similar figures are obtained when using data from the motor ROI. All statistics have been derived from data of *n* = 39 human participants who participated in one experiment. 1 TR = 1.25 s.

**Supplementary Figure S12:**
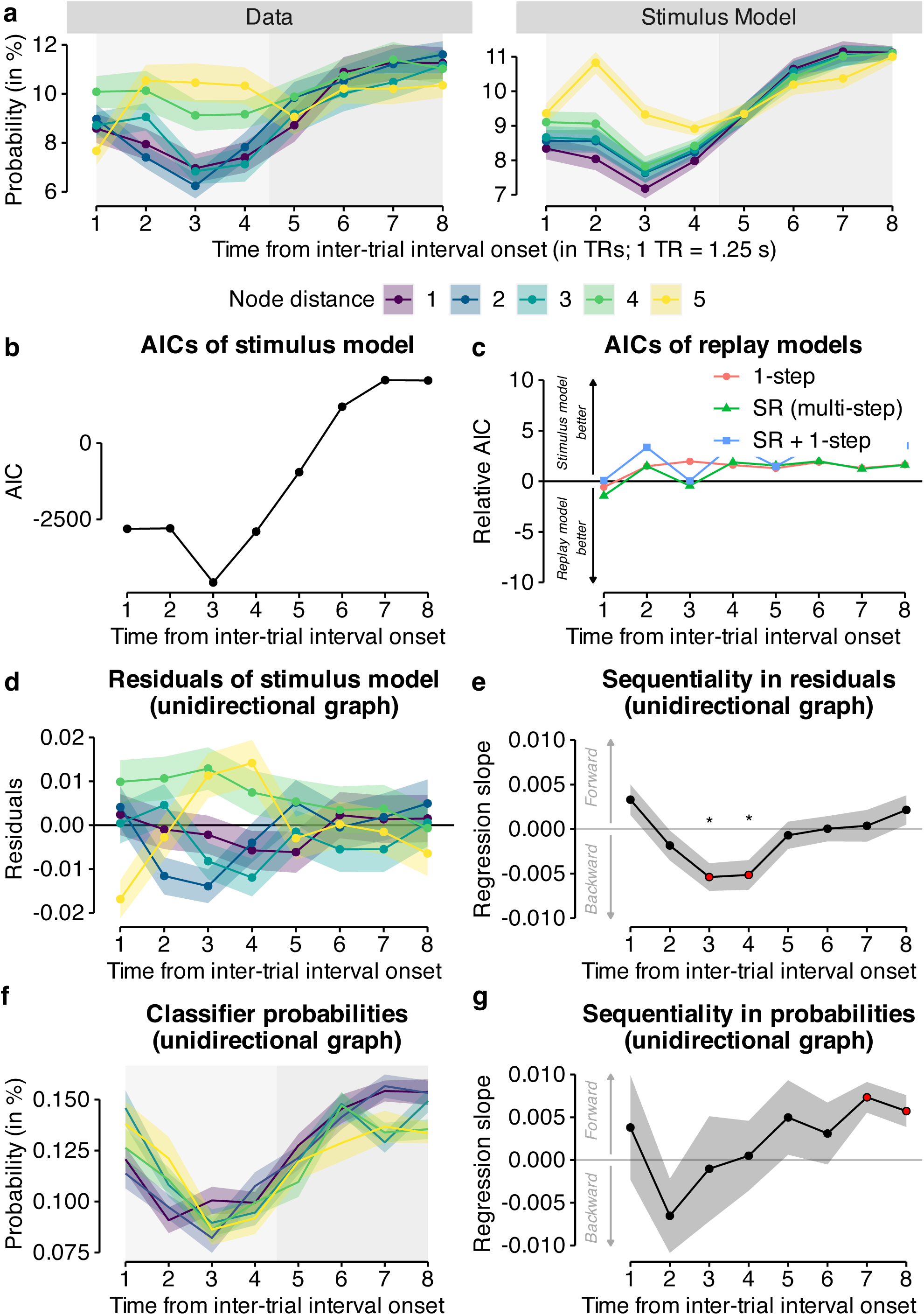
Classifier probabilities during on-task intervals in motor cortex. For details on each panel, see Fig. 5.

Task instructions in English

#### Box S1: Screen 1 of instructions for the training condition in session 1

Welcome to the study - Session 1!

Please read the following information carefully. If you have any questions, you can clarify them right away with the study instructor. Please lie as still and relaxed as possible for the
entire time.

Press any key to continue.

#### Box S2: Screen 1 of instructions for the training condition in session 1

Your task:

You are a zookeeper in training and have to make sure that all animals are in the right cages.

First you will learn in a training which animal belongs in which cage. We will now explain to you exactly how this task works.

Press any key to continue

#### Box S3: Screen 3 of instructions for the training condition in session 1

Training (Part 1)

You want to become a zookeeper and start your training today. First you will learn which animal belongs in which cage. You will see six cages at the bottom of the screen. Each of the six cages belongs to one of six animals. You will select a cage with the appropriate response key. Please keep your ring, middle and index fingers on the response keys the entire time so that you can answer as quickly and accurately as possible.

Press any key to continue

#### Box S4: Screen 4 of instructions for the training condition in session 1

During the training, the animals appear above their cages. Press the key for that cage as fast as you can and remember the cage where the animal belongs. Please press the correct button within 1 second. Please answer as quickly and accurately as possible. You will receive feedback if your answer was correct, incorrect or too slow. The correct cage will appear in green and the incorrect cage will appear in red.

Press any key to continue

#### Box S5: Screen 5 of instructions for the training condition in session 1

It is very important that you actively remember which animal belongs in which cage. You will get a higher bonus if you remember the correct assignment. The better you remember which animal belongs in which cage, the more money you earn! You will now complete one pass of this task, which will take approximately 2 minutes.

Press any key to continue.

#### Box S6: Screen 1 of instructions for the single trial condition in session 1

Training (part 2)

We will now check how well you have learned the assignment of the animals to their cages.

The animals will now appear in the center of the screen. You are asked to remember the correct cage for each animal, and then press the correct key as quickly as possible.

Press any key to continue.

#### Box S7: Screen 2 of instructions for the single trial condition in session 1

This time you respond only after the animal is shown. In each round, the animal will appear first in the center of the screen. Then please try to actively imagine the correct combination of animal, cage and response key. After that, a small cross will appear for a short moment. Then the cages appear and you can respond as quickly and accurately as possible. Please respond as soon as the cages appear, not earlier.

Press any key to continue.

#### Box S8

Screen 3 of instructions for the single trial condition in session 1 You have again 1 second to respond. Please respond again as fast and accurate as possible.

You will get feedback again if your response was wrong or too slow. If your response was correct, you will continue directly with the next round without feedback. You will now complete 8 passes of this task, each taking about 6 minutes. In between the rounds you will be given the opportunity to take a break.

Press any key to continue.

#### Box S9: Screen 1 of instructions for the single trial condition in session 2

Welcome to the study - Session 2!

We will check again if you can remember the assignment of the animals to their cages. The animals will appear in the center of the screen again. You are asked to remember again the correct cage for each animal and press the correct key as quickly as possible.

Press any key to continue.

#### Box S10: Screen 2 of instructions for the recall condition in session 2

You answer again only after the animal has been shown. In each round, the animal appears first in the center of the screen. Then please try to actively imagine the correct combination of animal, cage and answer key. After that, a small cross will first appear for a short moment.

Then the cages appear and you can answer as quickly and accurately as possible. Please respond as soon as the cages appear, not earlier.

Press any key to continue.

#### Box S11: Screen 3 of instructions for the single trial condition in session 2

You have again 1 second to respond. Please respond again as fast and accurate as possible.

You will get feedback again if your response was wrong or too slow. If your answer was correct, you will proceed directly to the next round without feedback. You will now complete a run-through of this task, which will again take approximately 6 minutes. After the round you will be given the opportunity to take a break. Press any key to continue.

#### Box S12: Screen 1 of instructions for the graph condition in session 2

You have finished the passage to memory! Well done! You are now welcome to take a short break and also close your eyes. Please continue to lie still and relaxed. When you are ready, you can continue with the instructions for the main task.

Press any key to continue.

#### Box S13: Screen 2 of instructions for the graph condition in session 2

Main task

Congratulations, you are now a trained zookeeper! Attention: Sometimes the animals break out of their cages! Your task is to bring the animals back to the right cages. When you see an animal on the screen, press the right button as fast as possible to bring the animal back to the right cage. This time you will not get any feedback if your answer was right or wrong. The more animals you put in the correct cages, the more bonus you get at the end of the trial!

The main task consists of 5 runs, each taking about 10 minutes to complete.

Press any key to continue.

#### Box S14: Screen 3 of instructions for the graph condition in session 2

You have again 1 second to respond. In the main task, you again respond immediately when you see an animal on the screen. Again, please respond as quickly and accurately as possible.

Between each round you will again see a cross for a moment. Sometimes the cross will be shown a little shorter and sometimes a little longer. It is best to stand by all the time to respond as quickly as possible to the next animal.

Press any key to continue.

#### Box S15: Screen 4 of instructions for the graph condition in session 2

Resting phases

After all the work as a zookeeper you also need rest. Before, between and after the main task we will take some measurements during which you should just lie still. During these rest periods, please keep your eyes open and look at a cross the entire time. Blinking briefly is perfectly fine. The background of the screen will be dark during the resting phases. Please continue to lie very still and relaxed and continue to try to move as little as possible. Please try to stay awake the entire time.

Please wait for the study instructor.

Task instructions in German

Removed from manuscript. According to bioRxiv policies, any information presented in a language other than English should be removed or substituted by the English translation

**Supplementary Figure S13:**
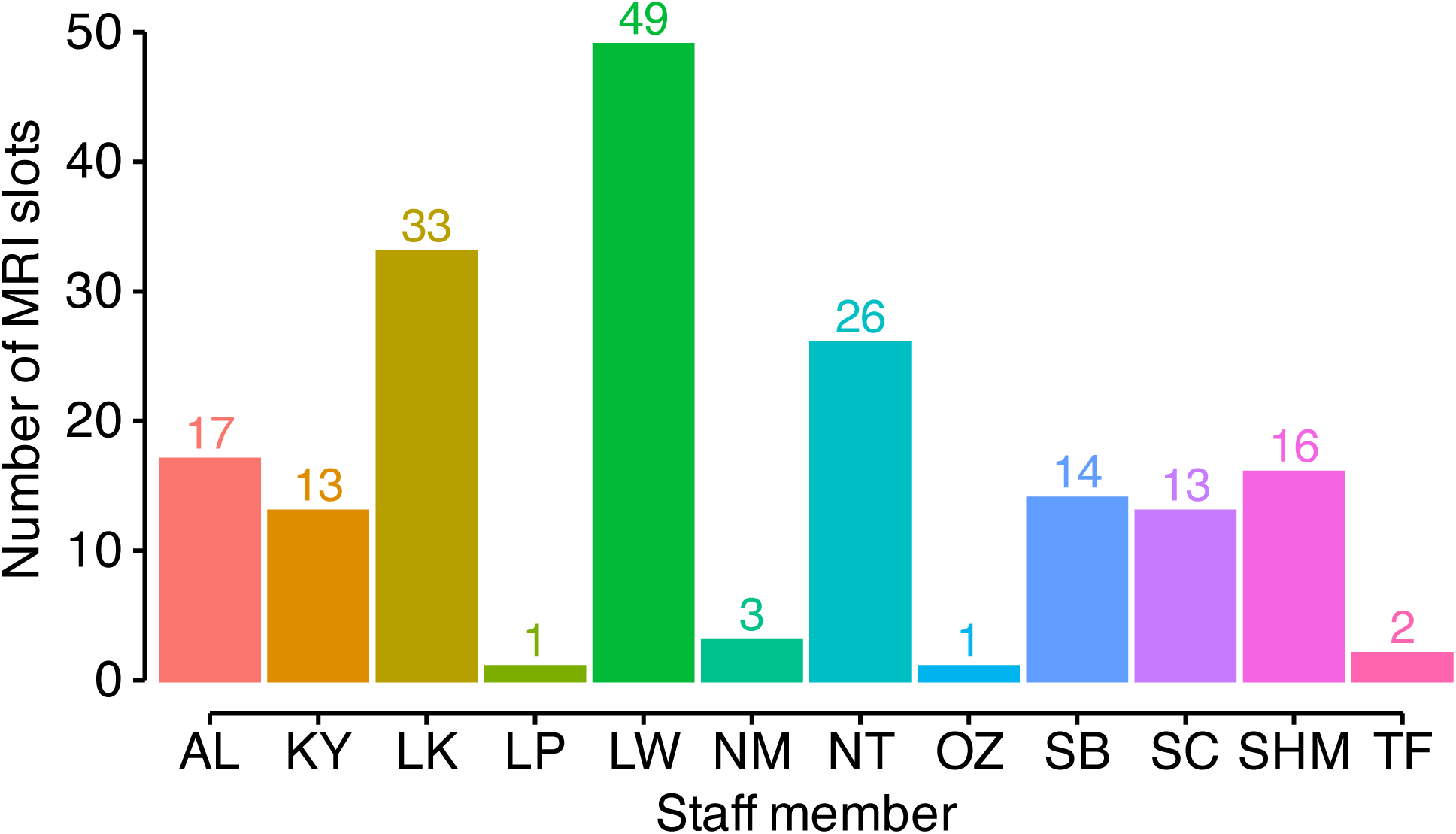
Real team effort during data collection. Number of MRI slots (y-axis and text labels) collected by each study staff member (x-axis; initials), indicating a a real team effort during data collection.

